# RBFOX2 deregulation promotes pancreatic cancer progression and metastasis through alternative splicing

**DOI:** 10.1101/2022.06.01.494386

**Authors:** Michelle Maurin, Mohammadreza Ranjouri, Katelyn Martin, Robert Miner, Justin Y. Newberg, Dongliang Du, Barbara Centeno, Jason B. Fleming, Xiaoqing Yu, Ernesto Guccione, Michael A. Black, Karen M. Mann

## Abstract

RNA splicing is an important biological process associated with cancer initiation and progression, yet in pancreatic cancer the role and regulation of splicing is not well understood. From a forward genetic screen in a mouse model of pancreatic cancer, we identified an enrichment of RNA binding proteins (RBPs) associated with the spliceosome. Here, we link deregulation of RBFOX2, an RBP of the FOX family, to pancreatic cancer progression and liver metastasis. We show that RBFOX2 regulation in pancreatic cancer occurs at both the RNA and protein level, and that nuclear localization of RBFOX2 is significantly reduced in poorly differentiated PDAC. Deregulation of RBFOX2 in PDAC is associated with an enrichment of exon exclusion events in transcripts encoding proteins involved in cytoskeletal remodeling and invadopodia programs that potentiate metastatic potential *in vivo*. Using splice-switching antisense oligonucleotides (AONs) and inducible cDNA isoforms, we demonstrate that RBFOX2 mediated exon exclusion in *ABI1* controls the abundance and localization of ABI1 protein isoforms in pancreatic cancer cells, and that ABI1 splice-switching enhances cellular phenotypes associated with cancer cell stemness. Together, our data identify a novel role for RBFOX2 deregulation in promoting PDAC progression through alternative splicing regulation.

## INTRODUCTION

Pancreatic ductal adenocarcinoma (PDAC) is a highly metastatic cancer driven by oncogenic *KRAS* mutations and inactivation of tumor suppressor genes *TP53, SMAD4* and *CDKN2A*. Importantly, these driver mutations are present in liver metastases [1, 2], the major site of disease dissemination and recurrence [3-5], suggesting that these events are necessary for disease maintenance. However, additional signaling, metabolic and regulatory processes are integral to disease progression. Recently, regulation of RNA splicing has gained attention for its importance in cancer development, and derived cancer splicing signatures have elucidated conserved events across cancer types [6, 7], the potential for splicing events to differentiate tumor tissue from normal [8-10], and the influence of alternative splicing events on therapy response.

RNA splicing is an integral biological process that regulates transcript stability and extends the repertoire of transcripts generated from a single genetic locus, contributing to a diversity of protein functions. Alternative splicing plays a major role in producing cell-type specific transcripts and is important for maintaining embryonic stem cell (ESC) pluripotency or promoting cellular differentiation [11-13]. Regulation of alternative splicing is complex, involving conserved protein complexes inclusive of RNA binding proteins that direct the specificity of splicing events [14]. In pancreatic cancer, individual splicing events have been linked to disease progression, patient survival and response to chemotherapy [15-19]. Analysis of global RNA splicing signatures in pancreatic cancer using TCGA RNA-seq data identified enriched pathways from alternatively spliced transcripts in metabolic processes, cell-cell adhesion and cytoskeletal organization, suggesting alternative splicing may impact processes important for cancer progression [16, 20]. A study by Wang *et al*. identified a gene set in PDAC regulated at multiple levels by mutation, differential gene expression and alternative splicing [21]. While the importance of alternative splicing in controlling cell fate is well established in development, the role and regulation of alternative splicing in cancer and the biological functions of protein isoforms generated from alternately spliced transcripts is not well understood.

Previously, we performed a forward genetic screen in mice using Sleeping Beauty insertional mutagenesis to identify genes and processes important for promoting pancreatic cancer progression [22, 23]. We identified an enrichment of genes encoding RNA binding proteins (RBPs), with a subset of these RBPs including *Mbnl1, Mbnl2* and *Rbfox2*, involved in alternative splicing. MBNL1, MBNL2 and RBFOX2 have known roles in promoting alternative splicing of exons in transcripts associated with ESC pluripotency and differentiation. RBFOX2 coordinately regulates alternative splicing with MBNL1 in the differentiation of iPSCs [12]. Further, RBFOX2 was identified as a key splicing regulator in ovarian and breast cancers responsible for the overlapping exon splicing patterns in these two cancers [9]. Both cancer types exhibit exon exclusion events associated with RBFOX2 downregulation, either by transcriptional control in ovarian cancer or by alternative splicing in breast cancer, resulting in a reduction of the nuclear RBFOX2 isoform. The role of RBFOX2 in directing alternative splicing of pancreatic cancer is not known.

In this study, we uncover deregulation of RBFOX2 as a driving event promoting pancreatic cancer progression and metastasis in mouse models. Utilizing *ex vivo* and *in vivo* approaches, we define RBFOX2-controlled alternative splicing events linked to cytoskeletal remodeling and specifically associate alternative splicing of *ABI1* to enhanced migratory and stem cell potential in pancreatic cancer. Our studies are the first to link RBFOX2-associated alternative splicing with pancreatic cancer progression.

## RESULTS

### *RBFOX2* is differentially expressed in pancreatic cancer

We identified an enrichment of progression driver genes encoding RNA binding proteins (GO: 0003723, *P*=1.2E-16, Enrichr) [24, 25] from our analysis of *Sleeping Beauty* (SB) insertional mutagenesis data from an *in vivo* forward genetic screen in a GEMM model of pancreatic cancer [22, 23]. A subset of these genes encodes proteins involved in the regulation of mRNA splicing via the spliceosome (GO: 0048024, *P*= 1.09E-09, Enrichr). An oncoprint of SB insertions in the eleven statistically defined splicing regulators (**Supplemental Figure 1, panel A**) shows the frequency and coincidence of insertions in these genes in a population of 172 pancreas tumors. *Mbnl1* and *Mbnl2*, the most frequent “hits” present in 25% and 31% of tumors, respectively, are RNA binding proteins previously described for their roles in alternative splicing in human ES cells [11-13]. *Rbfox2*, the third most frequently mutated gene (24% of tumors), encodes the RNA splicing factor RBFOX2, which is part of the FOX family of splicing factors and is the most ubiquitously expressed of the three family members, which also includes RBFOX1 and RBFOX3, primarily expressed in the brain and nervous system. Gene expression analysis of *MBNL1, MBNL2* and *RBFOX2* in normal pancreas and pancreatic tumors from GEO dataset GSE28735 revealed a significant decrease in expression of *MBNL2* and a significant increase in *RBFOX2* expression in tumors compared to normal (**Supplemental Figure 1, panels B** (FDR adj. *P*=0.0321) **and C** (FDR adj. *P*=0.006)) but no significant difference for *MBNL1* (**Supplemental Figure 1, panel D**). Further, we surveyed a panel of human cell lines derived from pancreatic ductal adenocarcinoma cancers (PDAC) for levels of total *MBNL2, MBNL1* transcripts by qPCR and found variable expression, with the highest levels in Panc1 and 8902 cells (**Supplemental Figure 1, panels E and F**). *RBFOX2* is also variably expressed across PDAC cell lines, with 4039 and Panc1 cells exhibiting the highest levels (**Figure 1, panel A**). *RBFOX2* expression has been linked to TGFß-driven epithelial-to-mesenchymal transition (EMT) in breast cancer [26-28]. PDAC cell lines 4039 and MiaPaCa2 are both mesenchymal-like, yet we found MiaPaCa2 cells exhibit half the amount of *RBFOX2* total transcripts as 4039 cells. Panc1 and PATC148 express both mesenchymal and epithelial markers, yet these two cell lines exhibit very different levels of *RBFOX2*, suggesting *RBFOX2* gene regulation is not solely dependent on EMT status in PDAC. RNA-seq gene expression signatures have been used to broadly sub-classify PDAC tumors as Classical (epithelial-like) or Basal (mesenchymal-like) [29-31]. To further investigate the relationship between *RBFOX2* expression and these two subtypes of PDAC, we interrogated the log2 *RBFOX2* expression in two RNA-seq datasets from resected human pancreatic cancer patients. We first sub-classified these datasets into Classical or Basal subtypes using defined gene expression signatures [32] (see Methods). From the E-MTAB-6830 RNA-seq dataset [33], we defined 56 classical tumors and 48 basal tumors and found a significant increase in *RBFOX2* in the basal subtype compared to the classical subtype (**Figure 1, panel B**, Wilcoxon t-test, *P*=0.00023). In contrast, *RBFOX2* expression was not significantly different between classical and basal subclasses from the RNA-seq data set EGAD00001004548 [1] (**Figure 1, panel C**, Wilcoxon t-test, *P*=0.0725). Together, these data suggest that *RBFOX2* gene expression levels are not consistently associated with specific PDAC subtypes.

**Figure 1.**
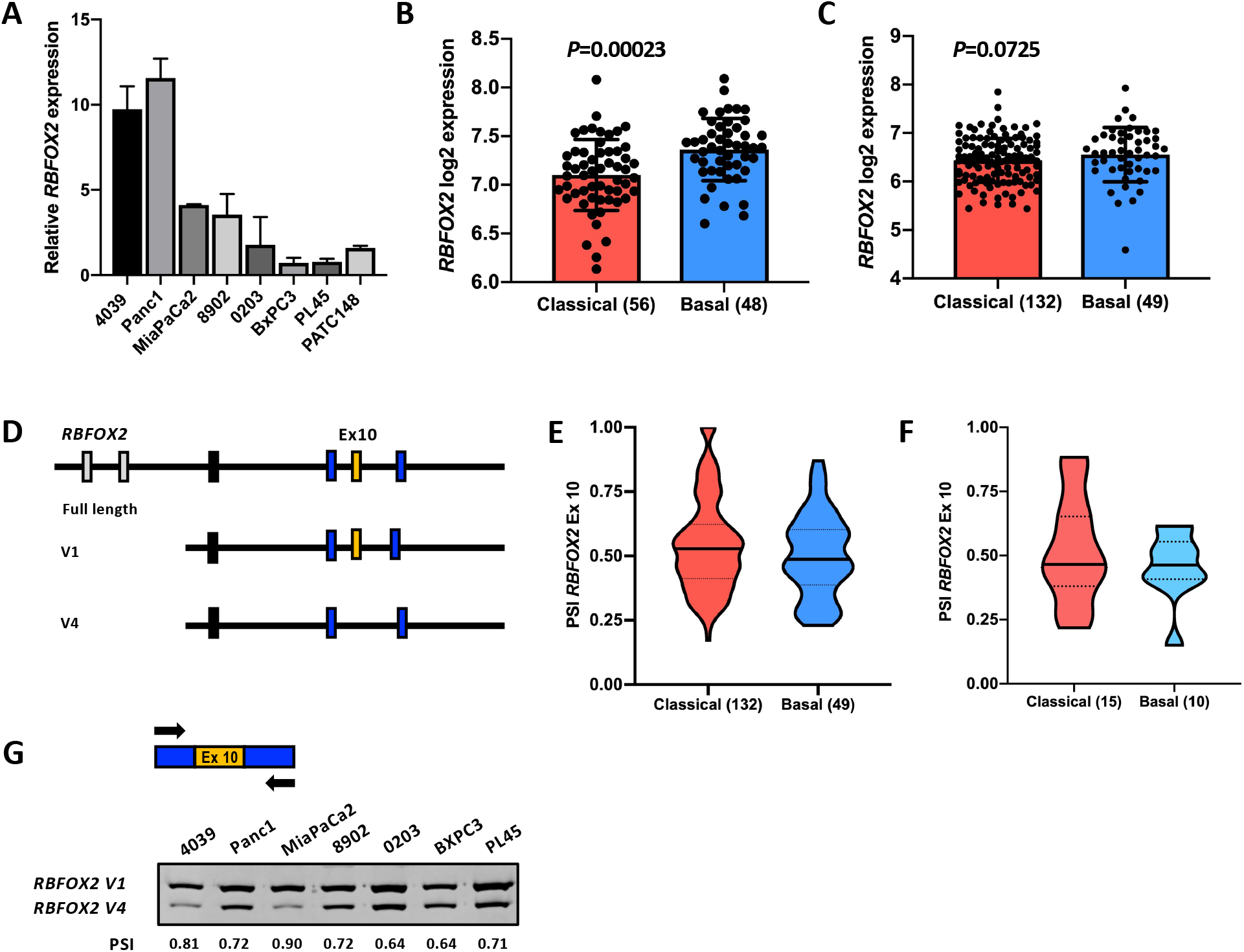
RBFOX2 is differentially expressed and alternatively spliced in pancreatic cancer. RBFOX2 is transcriptionally and post-transcriptionally regulated in pancreatic cancer. Quantitative PCR (qPCR) analysis of total *RBFOX2* using primers directed to the 3’ UTR (**A**) indicates that *RBFOX2* is most highly expressed in 4039 and Panc1 cells and most lowly expressed in PATC148 cells. qPCR analysis was performed over two independent cell line passages with technical replicates. *RBFOX2 gene* expression is higher in the basal subtype of human pancreatic cancer compared to the classical subtype in RNASeq data set E-MTAB-6830 (Wilcoxon *P*=0.00023, panel **B**). This distinction between PDAC subtypes and RBFOX2 expression is not significant in RNASeq data set EGAD00001004548 (Wilcoxon *P*=0.0725, panel **C**). *RBFOX2* is expressed as multiple RNA isoforms (**D**). Full length *RBFOX2* transcript (NM_001082578.2) contains two N-terminal exons not expressed in other isoforms, including v1 (NM_001031695.4) and v4 (NM_001082577.3) transcripts. Full length and v1 *RBFOX2* isoforms contain Exon 10 (ENSE00001578447) which is spliced out of the v4 transcript (NM_001082577). Analysis of RBFOX2 exon 10 (ENSG00000100320:015) inclusion in *RBFOX2* transcripts using RNASeq data (EGAD00001004548) shows a population mean percent spliced-in (PSI), a measure of exon inclusion, of 0.5 in resected pancreatic tumors for both the classical and basal subtypes (**E**). PSI is the percentage of transcripts including RBFOX2 exon 10 divided by the total RBFOX2 transcripts. The mean PSI for *RBFOX2* exon 10 in pancreatic cancer liver metastases is 0.45, with a tighter distribution in metastases with a basal gene signature compared to the classical gene signature (**F**). RT-PCR analysis using primers flanking *RBFOX2* exon 10 showed that exon 10 is predominantly included in *RBFOX2* transcripts across PDAC cell lines, with a lower PSI in epithelial-like lines (**G**).

### *RBFOX2* is alternatively spliced in PDAC

RBFOX2 is essential for human ESC cell survival [34] and is associated with developmental fates [12] and cancer progression [9]. *RBFOX2* transcripts expressed in PDAC are poorly characterized. The N-terminus of *RBFOX2* is alternatively spliced in a tissue-specific manner [35]. The first two exons of full-length *RBFOX2* transcripts (NM_001082578.2) encode a canonical nuclear localization signal (**Figure 1, panel D**) [35, 36]. *RBFOX2* transcripts can also utilize an internal start site and a nuclear localization signal provided by a 40-nucleotide, c-terminal exon [36]. This exon is alternatively spliced, and is present in the *RBFOX2* v1 transcript (NM_001031695.4) as exon 10 (ENSG00000100320:015). The v4 transcript (NM_001082577) lacks this exon, encoding an RBFOX2 protein lacking a nuclear localization signal. Interrogating the TCGA SpliceSeq database, which provides annotated alternative splicing events from TCGA RNA-seq datasets for over 30 cancer types [37] we confirmed alternative splicing of RBFOX2 exon 10 (exon 12 in SpliceSeq) in PDAC patients based on the range of the ratio of exon inclusion/exclusion in expressed transcripts, quantified as Percent Spliced-In (PSI, [38]). Further, a described autoregulatory alternative splicing event in *RBFOX2* exon 6 which removes part of the RNA recognition motif (RRM) [35] was not present in PDAC patient samples. Next, using an independent RNA-seq data set EGAD00001004548 from human pancreatic cancers, we aligned *RBFOX2* transcripts using DexSeq [39] and defined the percentage of transcripts containing exon 10 by calculating the PSI in both pancreas tumors and in liver metastases. We observed that the mean PSI for *RBFOX2* exon 10 is at or below 0.5 for both classical and basal subtypes of pancreatic cancer, with a large distribution of PSI values for individual tumors within the population illustrated by violin plots (**Figure 1, panel E**). The same observation was made for liver metastases, with a narrower distribution of PSI values for *RBFOX2*, more centered around 0.5 (**Figure 1, panel F**). These data suggest that on average, 50% of *RBFOX2* transcripts lack exon 10, which implies a reduction in nuclear RBFOX2. Further, we showed in a panel of PDAC cell lines with mesenchymal-like or epithelial-like characteristics that multiple RBFOX2 transcripts are expressed that either retain or lack exon 10, with an enrichment for transcripts containing exon 10 represented by the PSI >0.5 (**Figure 1, panel G**). Together, these data suggest that *RBFOX2* is regulated at both the transcriptional and post-transcriptional level in PDAC independent of the classical or basal subclasses of disease.

### Nuclear RBFOX2 protein expression is reduced in poorly differentiated PDAC

Given the complex regulation of *RBFOX2* at the RNA level in PDAC, we next investigated the abundance of RBFOX2 protein in in the panel of pancreatic cancer cell lines by Western blot analysis (**Figure 2, panel A**). We found that RBFOX2 protein is expressed as two isoforms, with higher expression of the 47 kD band in mesenchymal-like cells, compared to epithelial-like cells, as determined by vimentin (VIM) and E-cadherin (CDH1) expression, respectively. Panc0203 cells also exhibit high RBFOX2 protein levels, which may be due to amplification of the chromosomal region including the *RBFOX2* locus based on the Cancer Cell Line Encyclopedia (CCLE) copy-number variation data [40]. The PATC148 cell line, with low RBFOX2 abundance, exhibits both mesenchymal-like and epithelial-like properties and was derived from a PDX model of an invasive PDAC tumor that metastasized to the liver [41]. The upper 65 kD RBFOX2 band is poorly described and may represent a secondary modification. Epithelial-like PDAC cell lines exhibited lower abundance of this isoform compared to mesenchymal-like cells (**Figure 2, panel B**). Quantification of total RBFOX2 protein from three independent passages of each cell line is shown. Next, we investigated the abundance and distribution of RBFOX2 protein in resected pancreatic cancers. Using a tissue microarray constructed at Moffitt Cancer Center from resected pancreatic tumors, we found a significantly decreased percentage of RBFOX2-positive nuclei in tumor cells from PDAC patient samples diagnosed as moderate-poorly and poorly differentiated (n=22, average 57% percent positive nuclei) compared to well and well-to-moderately differentiated PDAC (n=31, average 68% percent positive nuclei) by immunohistochemistry, scoring an average of 1296 tumor nuclei per sample and excluding stroma (**Figure 2, panel C**, *P*=0.009, unpaired t-test). Representative images of tissue cores are shown for well-differentiated, moderately differentiated and poorly differentiated tumors (**Figure 2, panels D-F**). For a select few well-to-moderately differentiated PDAC samples, we observed 45% or fewer nuclei were positive for RBFOX2. These samples were diagnosed as invasive ductal adenocarcinoma, and representative cores are shown for invasive well-differentiated and moderately differentiated samples (**Figure 2, panels G and H**). Together, these data highlight the heterogeneity of nuclear RBFOX2 expression in PDAC, which correlates with the heterogeneity of *RBFOX2* transcripts encoding nuclear protein based on the PSI for exon 10. Further, these data associate reduced nuclear RBFOX2 with invasive and moderate-to-poorly differentiated pancreas cancers characterized by worse patient outcomes and increased liver metastasis [3, 4].

**Figure 2.**
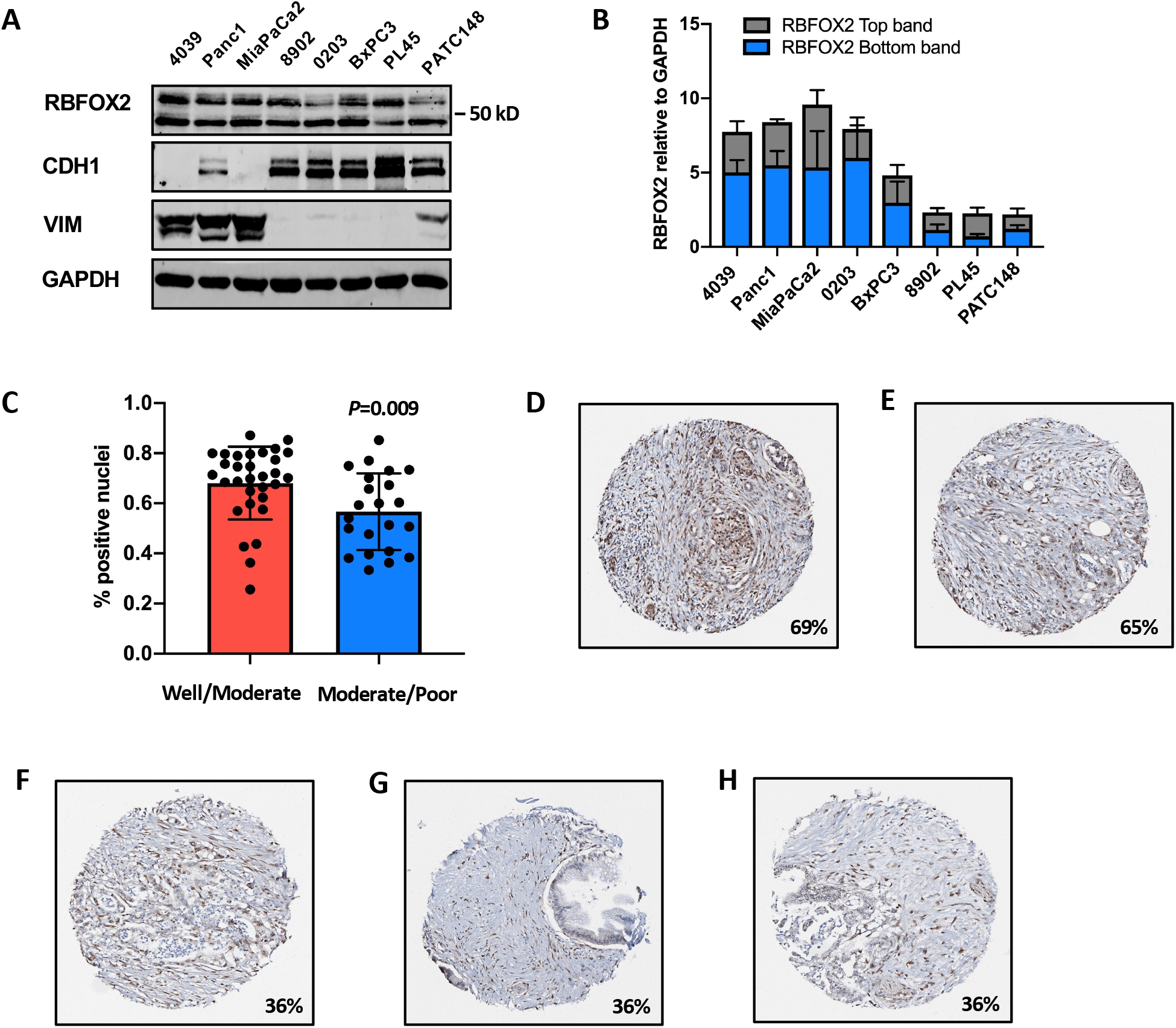
Nuclear RBFOX2 is decreased in poorly differentiated PDAC. Western blot analysis of RBFOX2 in a panel of human pancreatic cancer cell lines (**A**) revealed that RBFOX2 is present as two different molecular weights (65 kD and 47 kD) and is relatively more abundant in mesenchymal lines (marked by vimentin (VIM) expression). Relative quantitation of RBFOX2 demonstrates that PDAC cell lines preferentially express the 47 kD band (**B**). Analysis of RBFOX2 protein expression in 54 human pancreatic cancer samples using a tissue microarray from resected tumors constructed at Moffitt Cancer Center revealed that RBFOX2 nuclear abundance is significantly decreased in moderate-poorly differentiated PDAC (**C**, *P*=0.015, unpaired t-test) compared to well-differentiated PDAC. Representative images of RBFOX2 IHC from the TMA microarray for well-differentiated (**D**) moderately differentiated (**E**) and poorly differentiated PDAC sections (**F**) shows the distribution and intensity of RBFOX2 staining. Percent refers to the percent positive nuclei in each image. Well-differentiated (**G**) and moderately differentiated (**H**) diagnosed as invasive exhibit low nuclear positivity for RBFOX2.

### RBFOX2 depletion promotes aggressive phenotypes in pancreatic cancer cells

Given the association of decreased RBFOX2 in poorly differentiated PDAC, we wanted to further investigate the potential role of RBFOX2 in promoting pancreatic cancer progression. We performed stable depletion of RBFOX2 in MiaPaCa2, 4039 and Panc1 cells which expressed the highest level of protein. RBFOX2 protein levels were significantly depleted using a constitutively expressed shRNA (**Figure 3, panel A**) without changes in expression of VIM and CDH1, respectively. Quantification of the RBFOX2 protein knockdown is shown (**Figure 3, panel B**). We found no significant difference in the proliferative capacity of these isogenic pairs (**Figure 3, panel C**) grown under adherent conditions. However, when cells were plated under non-adherent conditions, we found that cells lacking RBFOX2 demonstrated a significantly increased number of live cells over a time course of 72 hours (*P*<0.0001, 2-factor ANOVA, **Figure 3, panel D**). To further characterize this phenotype, we interrogated the activation of cell survival pathways in Panc1 and MiaPaCa2 cells grown under non-adherent conditions for 48 hours and found that JNK phosphorylation is elevated in Panc1 and MiaPaCa2 cells depleted for RBFOX2 compared to RBFOX2-replete cells (**Supplemental Figure 2, panels A and B**), while AKT phospho-Ser473 was elevated in Panc1 cells depleted for RBFOX2 and unchanged in MiaPaCa2 cells (**Supplemental Figure 3, panels C and D**). Importantly, we found that RBFOX2 depletion significantly increased cell migration using a wound-healing assay shown for 4039 and Panc1 cells (**Figure 3, panels E and F**, 2-factor ANOVA, *P*<0.0001). As these cell lines express oncogenic mutant KRAS, an important driver in PDAC with known roles in promoting cellular proliferation and migration, we investigated whether RBFOX2 depletion influenced RAS activation by analysis of phospho-ERK. Interestingly, we showed in cells grown under adherent conditions that levels of phospho-ERK1 were depleted in Panc1 cells lacking RBFOX2 (**Supplemental Figure 3, panel E**). MiaPaCa2 cells do not express ERK1, while levels of phospho-ERK1/2 in 4039 cells were equivalent for the isogenic pair. Finally, RBFOX2 depletion significantly increased cell invasion, shown for 4039 cells (**Figure 3, panel G**, 2-factor ANOVA, *P*<0.0001) using an Incucyte chemotaxis assay. We confirmed our results using a second, doxycycline-inducible shRNA in MiaPaCa2, 4039 and Panc1 cell lines to efficiently deplete RBFOX2 (**Supplemental Figure 3, panels A and B**). We observed no change in cellular proliferative capacity upon RBFOX2 knockdown (**Supplemental Figure 3, panel C**). MiaPaCa2 cells exhibited significantly increased cell survival under non-adherent conditions (**Supplemental Figure 3, panel D**, *P*=0.0015, 2-factor ANOVA), while 4039 and Panc1 cells did not survive under non-adherent conditions with DOX treatment (data not shown). Induced RBFOX2 knockdown in 4039 cells significantly increased cell migration compared to control cells (**Supplemental Figure 3, panel E**, *P*<0.001, 2-factor ANOVA). Given that RBFOX2 nuclear depletion promotes aggressive *in vitro* phenotypes in PDAC cells, we next investigated whether RBFOX2 repletion could inhibit cell survival and migration *in vitro*. We stably transduced PATC148, PL45 and 8902 cells lines with doxycycline-inducible vectors expressing cDNAs for either the nuclear v1 isoform of RBFOX2 (i*RBFOX2* V1), the cytoplasmic v4 isoform (i*RBFOX2* V4) or a GFP control cDNA (i*GFP*). Western blot analysis confirmed induced expression of a 53kD Flag-tagged RBFOX2 v1 isoform (**Figure 4, panel A**) or a 53kD Flag-tagged RBFOX2 v4 isoform (**Supplemental Figure 4, panel A**). We achieved similar expression for each RBFOX2 isoform in each cell line panel. Quantification of total RBFOX2 in the v1 or v4 expressing lines compared to the GFP control is shown for PATC148 and 8902 cells (**Supplemental Figure 4, panel B**). Interestingly, induced expression of the RBFOX2 v4 cDNA led to a concomitant increase in endogenous RBFOX2, particularly for the 8902 isogenic pair (**Supplemental Figure 4, panel A**). Reconstituted expression of RBFOX2 did not change expression of VIM or CDH1. We confirmed that overexpression of RBFOX2 v1 did not alter levels of phospho-ERK1/2 (**Figure 4, panel B**). Nuclear-cytoplasmic cellular fractionation followed by western blot analysis confirmed the nuclear localization of the v1 isoform and exclusive cytoplasmic localization of the v4 isoform in PATC148 cells (**Supplemental Figure 4, panel C**). There was no change observed in cellular proliferative capacity for cells expressing v1 (**Figure 4, panel C**) or v4 (**Supplemental Figure 4, panel D**) from either PATC148 or 8902 cell lines compared to controls. PL45 cells were not very proliferative and were not characterized further. Induced v1 expression in PATC148 cells led to significantly reduced cellular survival under non-adherent conditions after 48 hours (**Figure 4, panel D**), while 8902 cells did not survive under non-adherent conditions. Finally, PATC148 cells with the v1 isoform showed significantly reduced cell migration by a wound-healing assay (**Figure 4, panel E**, *P*=0.0005, 2-factor ANOVA), while expression of v4 did not affect PATC148 cell migration (**Supplemental Figure 4, panel E**). Together, these data demonstrate that RBFOX2 depletion in PDAC cells promotes cellular migration, invasion and anoikis resistance, while nuclear RBFOX2 v1 reconstitution suppresses cell migration and anoikis resistance phenotypes in PATC148 cells with low endogenous RBFOX2 expression.

**Figure 3.**
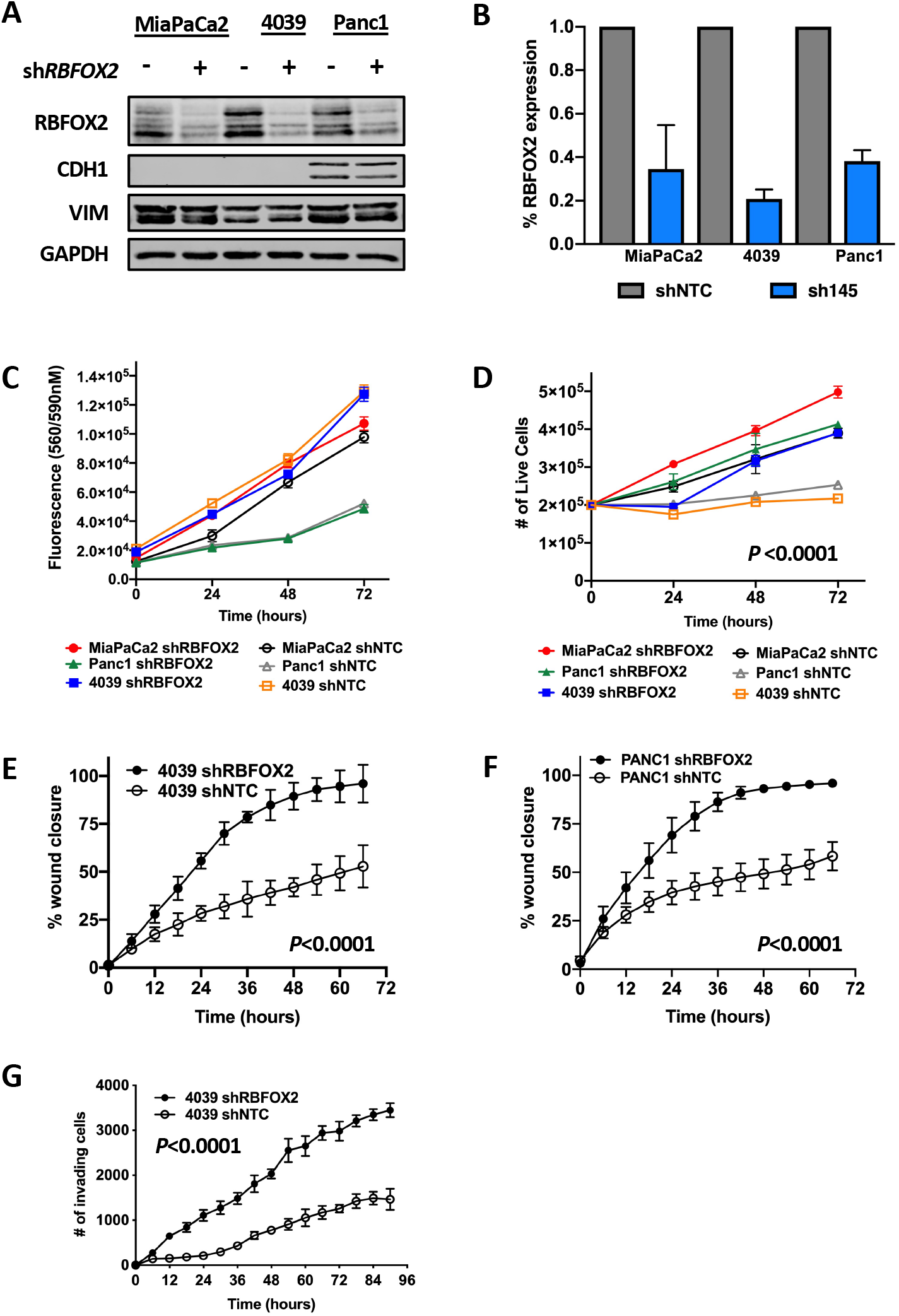
RBFOX2 depletion drives aggressive cellular phenotypes. ShRNA-mediated knock-down of RBFOX2 in mesenchymal-like PDAC cell lines MiaPaCa2, 4039 and Panc1 leads to a significant reduction in RBFOX2 protein (**A**) compared to cells stably transduced with a non-targeting shRNA (shNTC). No change in expression of epithelial marker CDH1 or mesenchymal marker vimentin (VIM) was observed with RBFOX2 knockdown. Cells with RBFOX2 depletion (sh*RBFOX2*) do not exhibit altered 2D cellular growth on plastic compared to control cells (shNTC) assessed using Cell Titer Blue assays (**C**). Cell survival under non-adherent conditions(**D**) is significantly increased in RBFOX2-depleted cells compared to control cells (*P*<0.0001, 2-factor ANOVA). Cell migration in 4039 (**E**) and Panc1 cells (**F**) is significantly increased upon RBFOX2 depletion using a wound healing assay (*P*<0.0001). Cell invasion is significantly increased upon RBFOX2 depletion in 4039 cells (**G**) (*P*<0.0001, 2-factor ANOVA).

**Figure 4.**
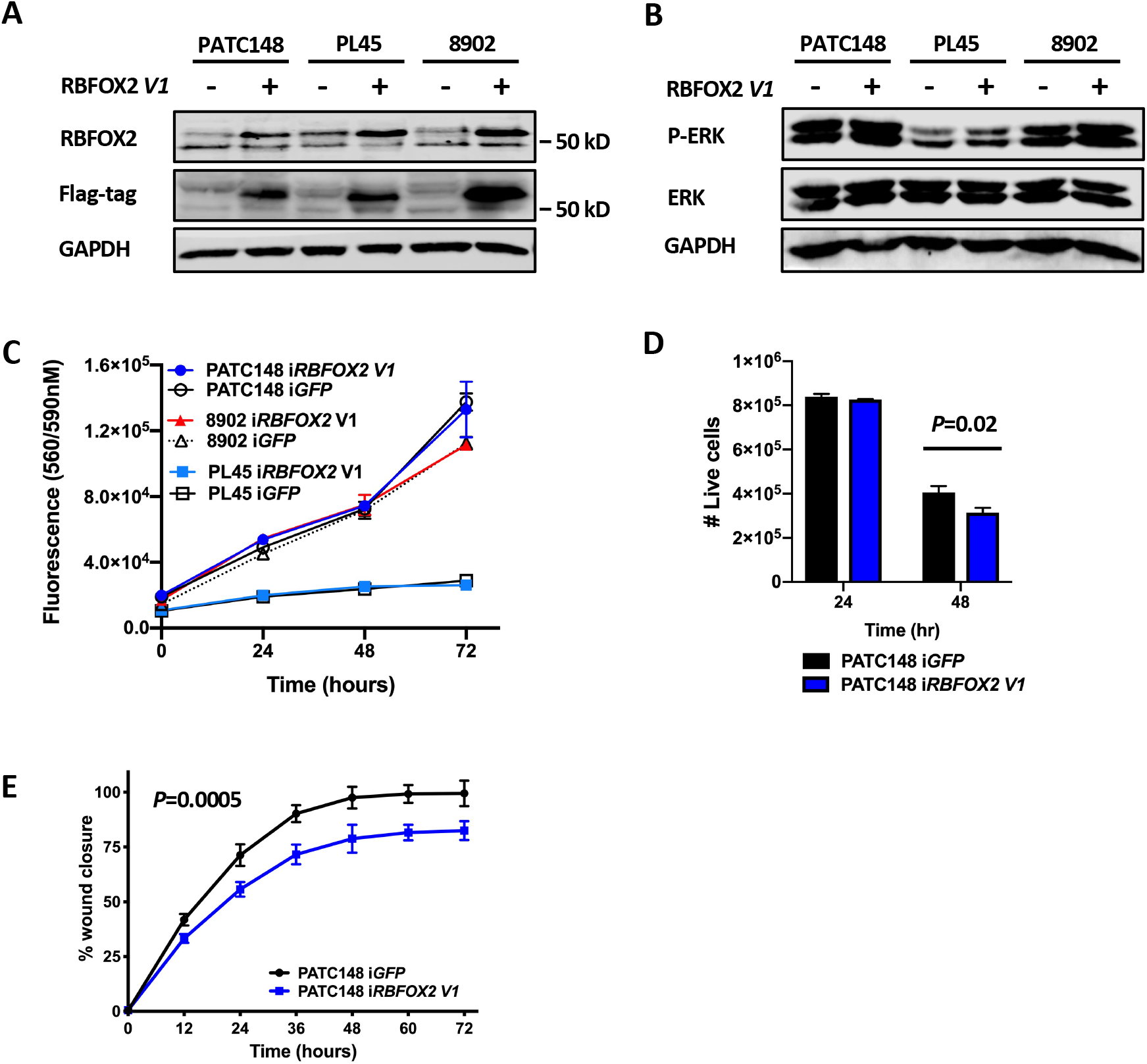
RBFOX2 repletion rescues cell migration in PDAC cells. Inducible *RBFOX2* with Flag-tagged v1 cDNA isoform expression reconstitutes RBFOX2 protein levels in PATC148, PL45 and 8902 cells (**A**). Detection of the Flag-tag confirms expression of the exogenous cDNA. Levels of total ERK or phospho-ERK1/2 (P-ERK) are unchanged across all isogenic pairs (**B**). No change in cellular growth was detected with RBFOX2 rescue (*iRBFOX2*) compared to control cells (*iGFP*) (**C**). Expression of RBFOX2 v1 (*iRBFOX2*) significantly reduced cell viability under non-adherent conditions compared to control cells (*iGFP*) in PATC148 cells (**D**, *P*=0.02, 48 hours). *iRBFOX2* significantly decreases cell migration in PATC148 cells (**E**, *P*=0.0005) compared to *iGFP* controls. *iRBFOX2* significantly decreases cell invasion in PATC148 cells using a wound-healing assay. Statistical analysis was performed using a 2-factor ANOVA.

### RBFOX2 depletion drives aggressive PDAC *in vivo*

We next investigated whether RBFOX2 depletion impacted PDAC tumor development *in vivo*. 4039 or Panc1 cells stably transduced with constitutive shRNA targeting *RBFOX2* or the non-targeting shRNA were introduced directly into the pancreas of NSG male and female mice by surgical injection. After 45 days, mice injected with 4039 cells with RBFOX2 depletion exhibited highly aggressive pancreatic disease, with large primary tumors and multiple lesions to the liver (**Figure 5, panel A**). Tumor volumes were increased in RBFOX2-depleted tumors compared to RBFOX2-replete tumors for 4039 cells (**Figure 5, panel B**, *P*=0.005, unpaired t-test) and Panc1 cells (**Supplemental Figure 5, panel A**, *P*=0.011). RBFOX2-depleted cells exhibited an enhanced ability to migrate and seed several organs based on the presence of foci expressing the GFP reporter. The incidence of metastasis to the liver was 80% for mice with RBFOX2-depleted tumors, defined by the presence of at least one lesion ≥1 mm in the liver, compared to only 10% of control mice (**Figure 5, panel C)**. Liver lesions from RBFOX2-depleted tumors exhibited both an increased size and multiplicity compared to RBFOX2-replete tumors (**Figure 5, panel D**). Notably, several mice with RBFOX2-depleted cells exhibited more liver lesions than could be measured. In addition, these mice exhibited foci in the mesentery, spleen and stomach. Panc1 cells with RBFOX2-depletion were also more migratory, with a 40% incidence of seeding to the liver and a 60% incidence of infiltration to the mesentery (**Supplemental Figure 5, panel B**). Again, both the size and multiplicity of metastatic lesions was increased in mice with RBFOX2-depleted tumors (**Supplemental Figure 5, panel C**). Histological analysis of pancreas tumors from RBFOX2 replete tumors by H&E staining showed tumor development with some normal pancreas within and surrounding the tumor (**Figure 5, panel D and Supplemental Figure 5, panel D**), while tumors derived from RBFOX2 depleted cells showed large areas of tumor growth with little stromal infiltrate and some normal pancreas surrounding the tumor area (**Figure 5, panel E and Supplemental Figure 5, panel E**).

**Figure 5.**
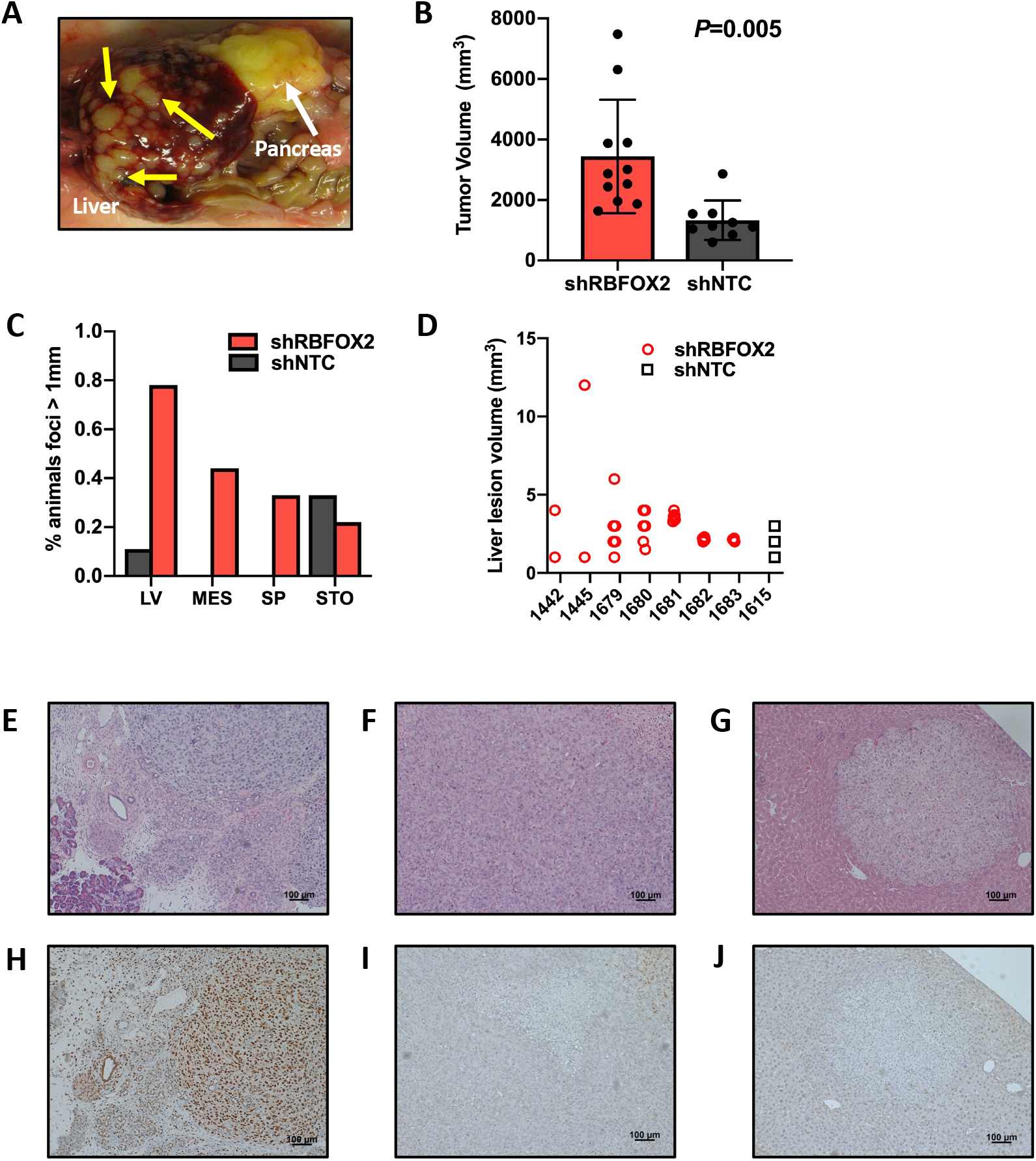
RBFOX2 depletion drives aggressive pancreatic cancer *in vivo*. Orthotopic injection of 4039 cells depleted for RBFOX2 promotes an aggressive disease in NSG mice (**A**). The pancreas tumor is labeled by the white arrow; multiple lesions in the liver are denoted by yellow arrows. Pancreas tumors derived from 4039 cells lacking RBFOX2 (red bars) have an increased tumor volume compared to tumors derived from control cells (black bars) (**B**, *P*=0.005, unpaired t-test). Mice with RBFOX2 depleted cells exhibit a higher incidence of metastatic lesions to the liver (LV), mesentery (MES), spleen (SP), stomach (STO), quantified as the percent of mice with at least one metastatic focus measured > 1 mm at necropsy (**C**). The volume of liver lesions (mm^3^) collected at necropsy is graphed on a per-animal basis (**D**). Only one mouse from the control cohort exhibited liver lesions. Histological analysis by H&E staining of pancreas tumors from 4039 cells expressing the non-targeting shRNA (**E**) shows tumor with surrounding acinar cells. Pancreas tumors with RBFOX2 depletion (**F**) have little surrounding normal pancreas. H&E staining from an RBFOX2-depleted liver lesion (**G**). Analysis of RBFOX2 expression by immunocytochemistry demonstrates robust RBFOX2 expression in tumors from replete cells (**H**) and an absence of signal in RBFOX2 depleted pancreas tumor (**I**) and resulting liver metastasis (**J**). RBFOX2 expression is decreased in the normal liver compared to normal pancreas.

Colonization of RBFOX2-depleted cells to the liver (**Figure 5, panel F and Supplemental Figure 5, panel F**) is surrounded by normal tissue. Immunohistochemistry confirmed RBFOX2 expression in the tumor and surrounding cells (**Figure 5, panel G and Supplemental Figure 5, panel G**) and the absence of RBFOX2 expression in the pancreas tumor (**Figure 5, panel H Supplemental Figure 5, panel H**) and liver lesions (**Figure 5, panel I and Supplemental Figure 5, panel I**). Note that the intensity of RBFOX2 staining in the normal pancreas is much higher than in the adjacent normal liver. We next investigated whether RBFOX2 repletion could decrease PDAC aggressiveness *in vivo*. We injected PATC148 cells with the DOX-inducible RBFOX2 v1 isoform or the inducible GFP control directly into the pancreas of NSG male and female mice by surgical injection. 7 days post-injection, mice were fed chow containing 200 ppm doxycycline to induce RBFOX2 v1 or the GFP control for the remainder of the study. After 45 days of growth from the time of injection, pancreas tumor volumes were not significantly different in the two cohorts (**Supplemental Figure 6, panel A**). However, the incidence of macroscopic lesions in surrounding organs, particularly in the mesentery, was reduced in mice injected with RBFOX2-replete cells (**Supplemental Figure 6, panel B**) compared to control. PATC148 cells do not metastasize to the liver. Histological analysis of RBFOX2-low (**Supplemental Figure 6, panel C**) and RBFOX2 replete pancreas tumors (**Supplemental Figure 6, panel D**) was very similar, with little normal pancreas observed. Immunochemistry to detect RBFOX2 in RBFOX2-low (**Supplemental Figure 6, panel E**) and RBFOX2 replete (**Supplemental Figure 6, panel F**) tumors showed heterogeneous expression, with slightly increased staining intensity for tumors with induced RBFOX2, suggesting that *in vivo* PATC148 cells maintain low RBFOX2 expression. Together, these *in vivo* data support a tumor suppressive role for RBFOX2 in controlling the metastatic potential of PDAC cells.

### RBFOX2 nuclear abundance affects alternative splicing of target exons

RBFOX2 is an RNA binding protein (RBP) that binds a highly conserved 6-nucleotide intronic recognition sequence (UGCAUG), recruits other splicing regulators, including HNRNPM [16], and facilitates alternative splicing of discrete exons [16-19]. RBFOX2 binding sites are highly conserved and RBFOX2 splicing regulation of target exons is tightly controlled by both the position and number of recognition sites in intronic sequences adjacent to spliced exons [34, 42]. RBFOX2 binding to a recognition site present downstream of an alternatively spliced exon facilitates the inclusion of the target exon in processed transcripts, while RBFOX2 binding to the recognition site present upstream of an alternatively spliced exon leads to exon skipping, due to RBFOX2-mediated inhibition of splicing factors to include the target exon in processed transcripts (**Figure 6, panel A**). RBFOX2 has been described as an important regulator of EMT splicing signatures in breast cancer models [28]. To define the profile of RBFOX2 splicing events in PDAC, we performed exon arrays in the isogenic cell line pairs replete and depleted for RBFOX2 (MiaPaCa2, 4039 and Panc1) and in the orthotopic primary tumors (4039 and Panc1) using Applied Biosystems Clariom D arrays (see Methods). We statistically identified 27 differentially spliced exons from a combined analysis of high-vs-low RBFOX2 cell lines and 43 differentially spliced exons from pancreas tumors replete and depleted for RBFOX2 (**Supplemental Table 1**). Eighteen of these alternatively spliced exons overlapped between the *in vitro* and *in vivo* analyses. In addition, we identified 56 intron retention, 14 alternative 5’ donor site and 4 alternative 3’ acceptor site events *in vitro* and 61 intron retention, 20 alternative 5’ donor site and 13 alternative 3’ acceptor site events. No intron retention or alternative 5’ donor site events were conserved between the *in vitro* and *in vivo* analyses, while only a single alternative 3’ acceptor site event was shared. In addition, we identified 332 differentially expressed protein coding genes with a 2-fold or greater change in expression between RBFOX2 replete and depleted samples that were not alternatively spliced (**Supplemental Table 2**). We focused on the alternative exon splicing events for further analysis. By overlaying mapped RBFOX2 binding sites [19] with these spliced exons, we confirmed that all 27 exons from the *in vitro* analysis and 35 of the exons from the *in vivo* analyses are flanked by conserved intronic RBFOX2 recognition sites present within 500 bp of the spliced exon, suggesting these exons are direct RBFOX2 targets. Further, the placement of the UCGAUG sequence corresponded with the direction of the exon splicing change upon RBFOX2 depletion (downstream for exon skipping and upstream for exon inclusion in the absence of RBFOX2). Focusing on this subset of non-overlapping 44 alternatively spliced exons, we found 11 map to transcripts encoded by statistically defined candidate driver genes in our SB_PDAC model, including *Abi1* and *Diaph1* (*Dia1*; see **Supplemental Figure 7** for oncoprint of insertions). Further, these 44 alternatively spliced exons are present in transcripts encoding structural or signaling proteins enriched in cytoskeletal function (GO:0005856, *P*=8.68E-07, Enrichr). The encoded proteins interact with a frequency greater than expected by chance, with multiple connections through Rac1 (**Figure 6, panel B**, STRING PPI *P* <7.48E-14) [43]. RBFOX2 depletion shifts the PSI of target exons in expressed transcripts. We validated exon splicing events in our isogenic cell line pairs replete and depleted for RBFOX2 by RT-PCR using primers flanking the target exon and further refined the list of differentially spliced RBFOX2 target exons based on a 20% change in PSI with RBFOX2 depletion. In RBFOX2-replete cells, spliced transcripts containing the RBFOX2 target exon were predominant, as illustrated by *DIAPH1*. For other genes, including *ABI1* and *ECT2*, multiple transcripts with inclusion or exclusion of the RBFOX2 target exon were co-expressed (**Figure 6, panel C**). *ABI1, DIAPH1, DIAPH2* and *ECT2* showed consistent exon skipping across cell lines with constitutive (**Figure 6, panel C**) or induced RBFOX2 depletion (**Supplemental Figure 8, panel A**), with similar reduction of RBFOX2 protein determined by western blot (**Supplemental Figure 8, panel B**). In MiaPaCa2 cells, skipping of the RBFOX2-target exon in *DIAPH2* occurs in the presence of RBFOX2, suggesting that in this cell line *DIAPH2* exon splicing is regulated by other RBP(s). Nuclear RBFOX2 repletion in PATC148, 8902 and PL45 cells reverted exon skipping and increased the PSI for *ABI1, DIAPH2* and *ECT2*, confirming these exons are direct RBFOX2 targets in PDAC cells. Across all cell lines, the majority of *DIAPH1* transcripts retained the RBFOX2 target exon, which was skipped upon RBFOX2 depletion. *DIAPH1* splicing was unaffected by RBFOX2 repletion (**Figure 6, panel D**). As expected, repletion of the cytoplasmic RBFOX2 v4 isoform did not revert RBFOX2-mediated splicing of these targets (**Supplemental Figure 8, panel C**). The PSI for *ECT2* was less than 0.5 in all control cells, suggesting that the RBFOX2 target exon is regulated in part by other splicing factors. Retention of this exon is clearly regulated by RBFOX2 based on our depletion and repletion experiments. Because the PSI for *ABI1* in 8902 control cells was similar to that achieved by RBFOX2 depletion in the mesenchymal-like cell lines, we further investigated the RBFOX2 splicing profile in 8902 cells. We generated isogenic cell line pairs with RBFOX2 knockdown using either the constitutive or inducible RBFOX2 shRNAs with similar reduction of RBFOX2 protein determined by western blot (**Supplemental Figure 8, panel D**). Reduction of RBFOX2 in 8902 cells further reduced the PSI for *ABI* and *ECT2* (**Supplemental Figure 8, panel E**) and facilitated exon exclusion in *DIAPH1* and *DIAPH2*. These data suggest that alternative splicing of these RBFOX2 target exons is dependent on RBFOX2 abundance and is not dependent on the epithelial-like or mesenchymal-like properties of PDAC cells. In contrast, alternative splicing of the RBFOX2 target exons in *APBB2, CTTN, UAP1* or *UTRN* transcripts does appear to be influenced by the mesenchymal properties of PDAC cells, as the PSI in these cell lines was reduced by at least 50% in MiaPaCa2 and 4039 cells and reduced by less than 30% or was unchanged for Panc1 and 8902 cells with RBFOX2 depletion. For *MYO18A*, exon skipping appears to be associated with epithelial-like properties, as the control PSI for 8902 cells was reflective of the PSI achieved by RBFOX2 reduction in MiaPaCa2, 4039 and Panc1 cells (**Supplemental Figure 8, panel F**). RBFOX2 loss increases the PSI for *ADD3, EXOC1*, and *MAP3K7* for all cell lines to varying degrees. Next, we confirmed using TCGA SpliceSeq [37] a public database of annotated splicing events across 33 cancer types from TCGA, that the majority of the RBFOX2 target exons we identified are alternatively spliced in pancreatic cancer. Further, we identified a positive correlation between *RBFOX2* gene expression (log2 Transcripts Per Million (TPM)) and the PSI for 42 of the RBFOX2 target exons with exact matches for exon coordinates from RNA-seq dataset EGAD00001004548 (Pearson correlation *P*=0.0009) using a permutation test to compare the average absolute correlation between the PSI of spliced exons and *RBFOX2* expression with the average absolute correlation of randomly selected exons and *RBFOX2* expression in resected human PDAC patients (see Methods and histogram **Supplemental Figure 9, panel A** and **Supplemental Table 3**). Individual correlation plots for RBFOX2 target exons with *RBFOX2* expression are shown for *DIAPH1, ABI1* and *DIAPH2* (**Figure 6, panels E-F**). We conducted a similar analysis to define a positive correlation between *RBFOX2* gene expression (log2 TPM) and the PSI of 42 RBFOX2 target exons in liver metastases (Pearson correlation *P*=0.0116). Individual correlation plots for RBFOX2 target exons with *RBFOX2* expression for *DIAPH1, ABI1* and *DIAPH2* in liver metastases are shown (**Supplemental Figure 9, panels B-D** and **Supplemental Table 3**). Finally, we investigated the population distribution of the RBFOX2 target exon PSIs in pancreas tumors (**Figure 6, panel H**) and confirmed the majority of transcripts exclude RBFOX2 target exons, with a mean PSI of 0.35 for *ABI1* and ≤0.25 for *DIAPH2* and *ECT2* transcripts. Similar trends were observed in liver metastases (**Figure 6, panel I**). Together, these data demonstrate that in pancreatic cancer, RBFOX2 controls exon splicing events in transcripts encoding proteins with important roles in cancer progression.

**Figure 6.**
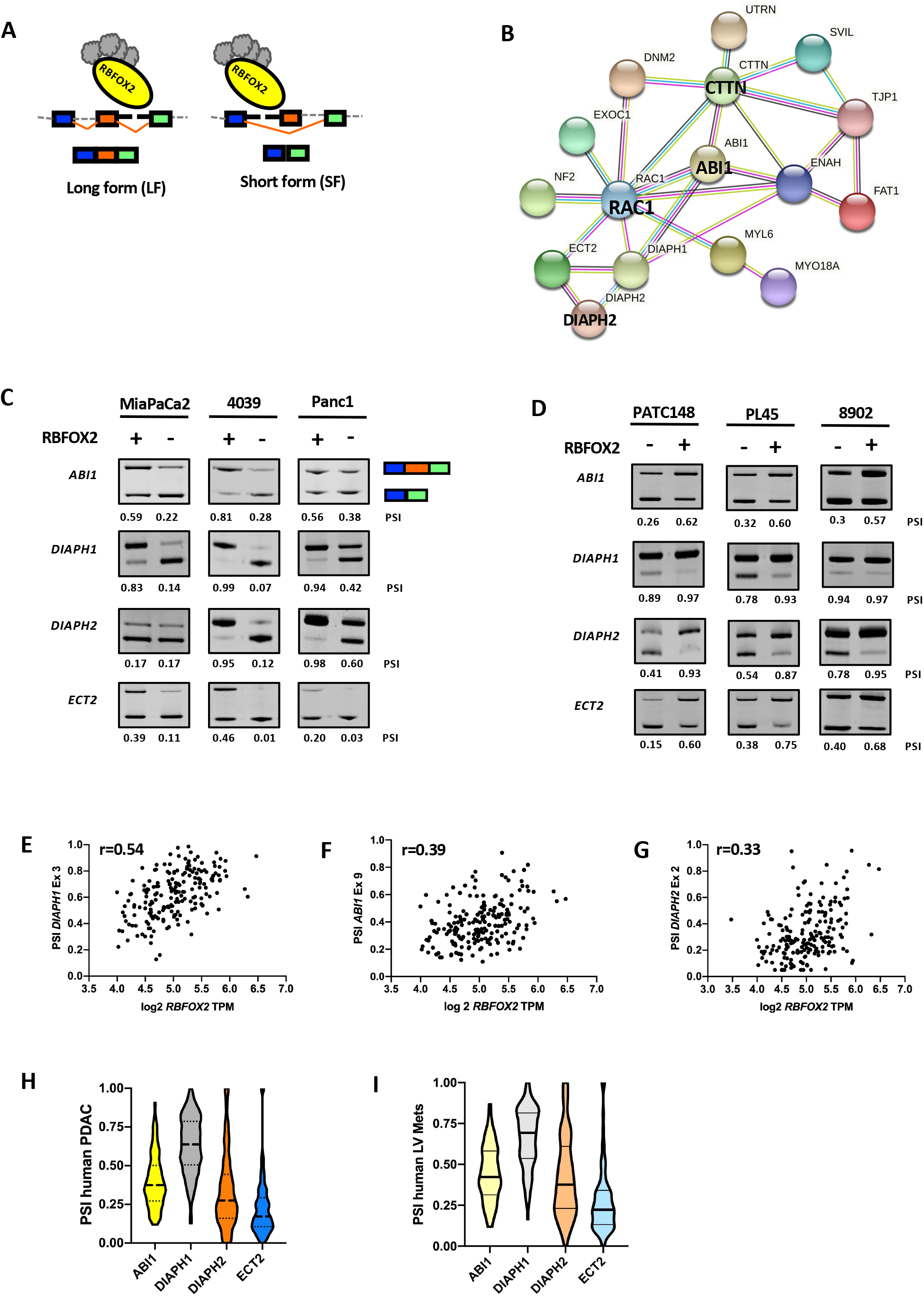
RBFOX2 depletion promotes exon skipping in invadopodia regulators. RBFOX2 binds a 6-nucleotide intronic recognition sequence (UGCAUG) and recruits splicing machinery to facilitate splicing of target exons (**A**). RBFOX2 binding downstream of the target exon (orange) facilitates its inclusion, generating the long form (LF) of the processed mRNA. RBFOX2 binding upstream of the target exon facilitates its exclusion, generating the short form (SF) of the processed mRNA. (**B**) RBFOX2 splicing targets are significantly enriched for protein interactions through RAC1 (STRING PPI *P* <7.48E-14). (**C**) RT-PCR analysis of RBFOX2 target exons in *ABI1, DIAPH1, DIAPH2* and *ECT2* shows a shift towards exon exclusion in the absence of RBFOX2 in 4039, Panc1 and MiaPaca2 isogenic cell line pairs. PSI values (Percent spliced-in) calculated for each isogenic pair quantify the decrease in transcripts encoding the long isoform (LF) upon RBFOX2 depletion. Reconstitution of RBFOX2 in PATC148, 8902 and PL45 cells promotes exon inclusion of RBFOX2 target exons in *ABI1, DIAPH1, DIAPH2* and *ECT2* (**D**). PSI values quantify the increase in transcripts encoding the long mRNA isoform (LF). RBFOX2 target exon abundance (calculated as PSI) positively correlates with *RBFOX2* expression (Log2 TPM) using Pearson’s statistical correlation in RNASeq data set EGAD00001004548 for *DIAPH1* (**E**, r=0.54), *ABI1* (**F**, r=0.39) and *DIAPH2* (**G**, r=0.33). Violin plots for the population PSI for RBFOX2 target exons shows exon skipping is enriched for *ABI1, DIAPH2* and *ECT2* in pancreas (**H**) and liver (**I**) but not for *DIAPH1*.

### RBFOX2 depletion promotes invadopodia formation and ABI1 redistribution through *ABI1* alternative splicing

RBFOX2 spliced target exons in PDAC are enriched in transcripts involved in cytoskeletal remodeling and cell adhesion, including *ABI1*. ABI1 is part of the WAVE regulatory complex (WRC) present at the leading edge of cells that regulates actin remodeling through the interactions with the Arp2/3 complex to promote lamellopodia formation. The WRC is activated through ERK phosphorylation of WAVE and ABI1 [29] and by Rac1-induced phosphorylation of Sra1 [30]. To determine whether RBFOX2 loss promotes increased lamellopodia formation, we grew RBFOX2 replete and depleted cells on gelatin-coated coverslips and stained them for phalloidin to mark F-actin. We observed increased membrane protrusions upon RBFOX2 depletion, particularly for 4039 cells (**Figure 7, panel A**). We quantified the number of F-actin foci in RBFOX2 replete and depleted cells using the ComDet plugin for ImageJ (see methods) and found a significant increase in the number of foci per cell upon RBFOX2 depletion in 4039 and MiaPaCa2 cells, with an average of 25 cells scored per condition (**Figure 7, panel B**). Rac1 activation is well known to play an important role in cytoskeletal remodeling through binding of WAVE regulatory components [44, 45]. Notably, we found no difference in levels of GTP-bound Rac1 in RBFOX2 replete and depleted cells using GTP-pulldown assays (**Figure 7, panel C**), suggesting the phenotypes we observe upon RBFOX2 depletion are downstream of Rac1 activation. Importantly, ABI1 exhibits a shift in protein isoform abundance in response to RBFOX2 depletion, consistent with the splice shift we observe in *ABI1* transcripts (**Figure 7, panel D**). Interestingly, we found that PDAC cell lines exhibit different profiles of ABI1 isoforms. RBFOX2-replete MiaPaCa2 cells exhibited the greatest abundance of ABI1 and co-expressed ABI1 full length (NP_001171590.1) and ΔEx9 (NP_001171591.1) protein isoforms. Upon RBFOX2 depletion, we observed a distinct shift to predominantly express ABI1ΔEx9. 4039 cells replete for RBFOX2 expressed ABI1 full length exclusively but exhibited an increase in ABI1ΔEx9 with RBFOX2 depletion. Panc1 cells exhibit three ABI1 isoforms, with ABI1 full length the predominant isoform in the presence of RBFOX2 and a lower predominant isoform in the absence of RBFOX2. Finally, PATC148 cells also exhibit 3 ABI1 isoforms, while RBFOX2 repletion results in an enrichment of ABI1 FL protein, consistent with increased exon inclusion (**Figure 7, panel E**). These data demonstrate that the abundance of ABI1 protein isoforms can be predicted by the PSI of exon 9 for *ABI1* RNA transcripts. To further investigate the localization of ABI1 in RBFOX2-depleted cell lines, we performed siRNA-mediated *RBFOX2* knockdown in Panc1 and MiaPaCa2 cells (**Supplemental Figure 10, panel A**) and confirmed isoform switching of ABI1 at both the RNA and protein level (**Supplemental Figure 10, panels B and C**). We then stained cells for ABI1 and phalloidin. MiaPaCa2 cells replete for RBFOX2 exhibited diffuse staining for ABI1 and F-actin (**Figure 7, panel F**). In contrast, MiaPaCa2 cells with RBFOX2 depletion exhibited ruffled edges and a redistribution of F-actin to the cell periphery with a coincident redistribution of ABI1 foci to the cell surface. Similar results were observed in Panc1 cells (**Supplemental Figure 10, panel D**). These data suggest RBFOX2 depletion promotes cellular remodeling associated with both enrichment and cellular redistribution of the ABI1ΔEx9 isoform. To further characterize the function of ABI1 isoforms in PDAC cells, we generated isogenic cell lines expressing endogenous ABI1 and RBFOX2 levels with inducible tagged cDNAs for either *ABI1*ΔEx9 (NM_001178120.2) or *ABI1* full-length (NM_001178119.2). *ABI1* exon 9 splicing analysis by RT-PCR confirmed induced expression of the *ABI1*ΔEx9 cDNA promoted a decrease in the abundance of the endogenous long-form transcript (ΔPSI 0.56 to 0.39) in Panc1 cells, while induction of ABI1 full-length shifted the proportion of the ABI1 isoforms to the long-form almost exclusively (ΔPSI 0.24 to 0.93) (**Figure 8, panel A**). Western blot analysis confirmed increased expression of ABI1ΔEx9 in Panc1 cells but showed little change in expression of ABI1 full-length. Notably, the total amount of ABI1 protein in Panc1 cells with induced *ABI1*ΔEx9 was increased. In PATC148 cells, we observed a dramatic shift in expression of ABI1 full-length compared to control empty vector (**Figure 8, panel B**). We then performed immunocytochemistry to look at the localization of ABI1 in PATC148 cells with induced ABI1 full length or inducible control vector in the presence of low endogenous RBFOX2 (shown in **Figure 4, panel A**). Similar to our observations in cells with forced depletion of RBFOX2, we observed that the PATC148 cells with low endogenous RBFOX2 show elongated projections with F-actin at the periphery and overlapping ABI1 localization, with some observed in the cell body (**Figure 8, panel C**). In contrast, induced expression of ABI1 FL resulted in a distinct cellular phenotype denoted by an absence of cellular projections and concentrated staining of ABI1 and F-actin surrounding the nucleus. This observation is consistent with the cellular morphology of Panc1 cells expressing endogenous ABI1 FL and suggests the presence of the ABI1ΔEx9 splice form has an important role in promoting elongation of F-actin filaments and lamellopodia in pancreatic cancer cells. Next, we showed that induction of ABI1ΔEx9 in Panc1 cells significantly increased cell migration compared to induced control cells (**Figure 8, panel D**, *P*<0.0001 2-factor ANOVA). To examine the contribution of endogenous ABI1 FL to cell migration in the context of induced ABI1ΔEx9, we performed siRNA-mediated knockdown of ABI1 targeting the 3’UTR of endogenous ABI1 (**Figure 8, panel E**), achieving 50% knockdown at the protein level which resulted in significantly decreased cellular migration in Panc1 cells. We found induction of *ABI1ΔEx9* rescued the decreased cellular migration observed with knockdown of endogenous ABI1 (**Figure 8, panel F**), demonstrating that ABI1ΔEx9 promotes cellular migration.

**Figure 7.**
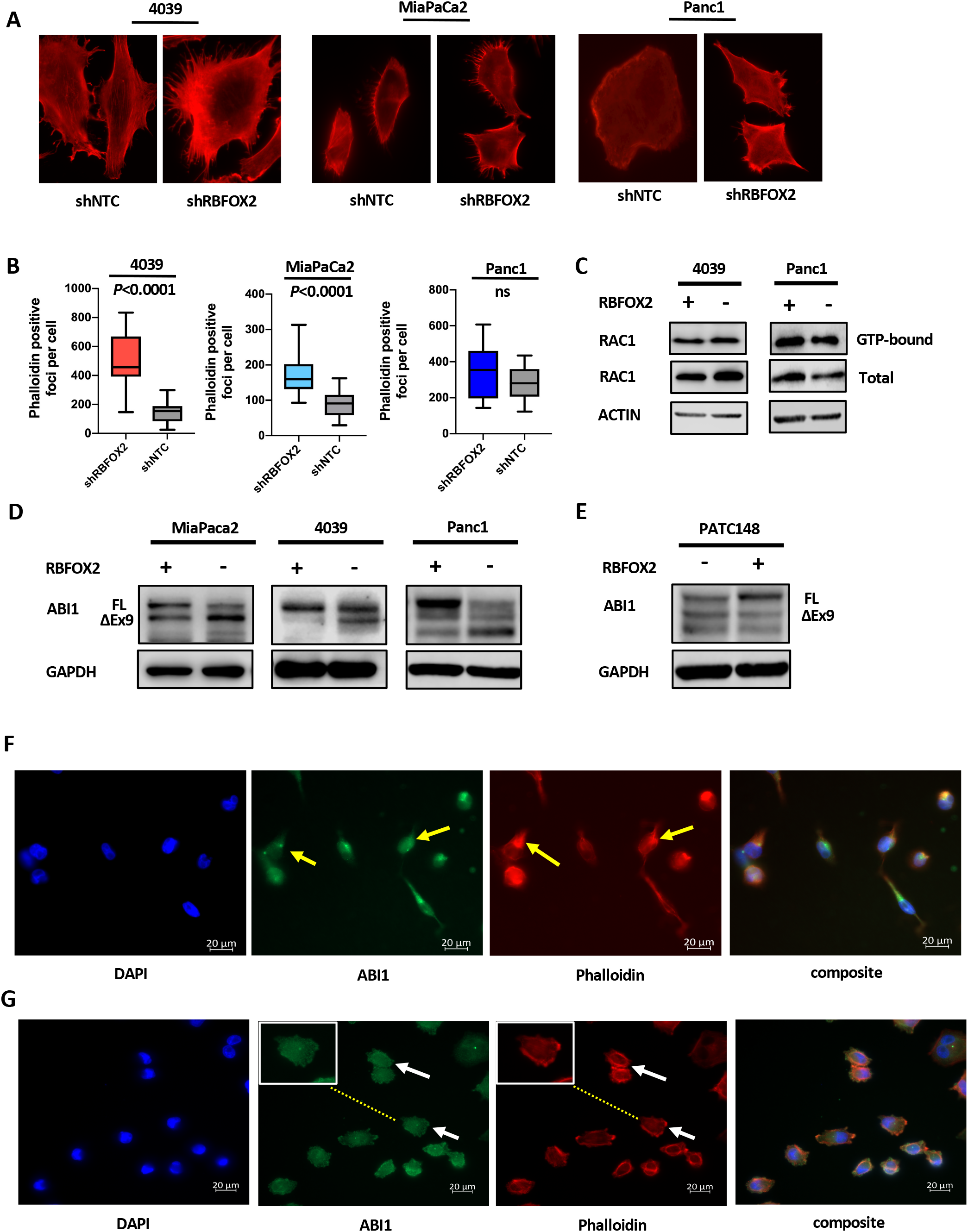
RBFOX2-mediated spliceforms of Rho GTP-regulated proteins promotes invadopodia formation in PDAC cells. 4039 and MiaPaCa2 cells with RBFOX2 depletion demonstrate an increase in the number of phalloidin positive projections compared to control cells (**A**). Panc1 cells exhibit morphological changes but few projections. 4039 and MiaPaCa2 cells depleted for RBFOX2 show a significant increase in phalloidin positive foci (**B**, *P*< 0.0001, unpaired t-test). The number of phalloidin positive foci in Panc1 cells replete and depleted for RBFOX2 is not significantly different. An average of 25 cells were scored for each cell line. Levels of GTP-bound Rac1 in 4039 and Panc1 cells replete and depleted for Rac1 were unchanged in cells repleted for RBFOX2 compared to controls using a GTP pulldown assay (**C**). Depletion of RBFOX2 promotes a shift in protein abundance of ABI1 isoforms towards the ΔEx9 isoform for MiaPaca2 and 4039 cells. ABI1 full-length is reduced in Panc1 in RBFOX2-depleted cells and a smaller ABI1 protein isoform is increased lacking ΔEx9 and neighboring exons (**D**). RBFOX2 repletion in PATC148 cells promotes a shift towards increased abundance of the ABI1 full-length isoform (**E**). ABI1 isoforms show differential cellular localization in RBFOX2 replete and depleted cells. In RBFOX2 replete cells (siNTC), ABI1 and F-actin are distributed and overlap in the cytoplasm and the perinuclear regions (**F**, yellow arrowheads). siRNA-mediated RBFOX2 depletion in MiaPaCa2 cells recapitulates morphological changes and increased cellular projections observed with shRNA-mediated RBFOX2 knockdown. ABI1 and F-actin are redistributed to the cellular periphery and are present in cellular projections in RBFOX2 depleted cells (**G**, white arrowheads and inset).

**Figure 8.**
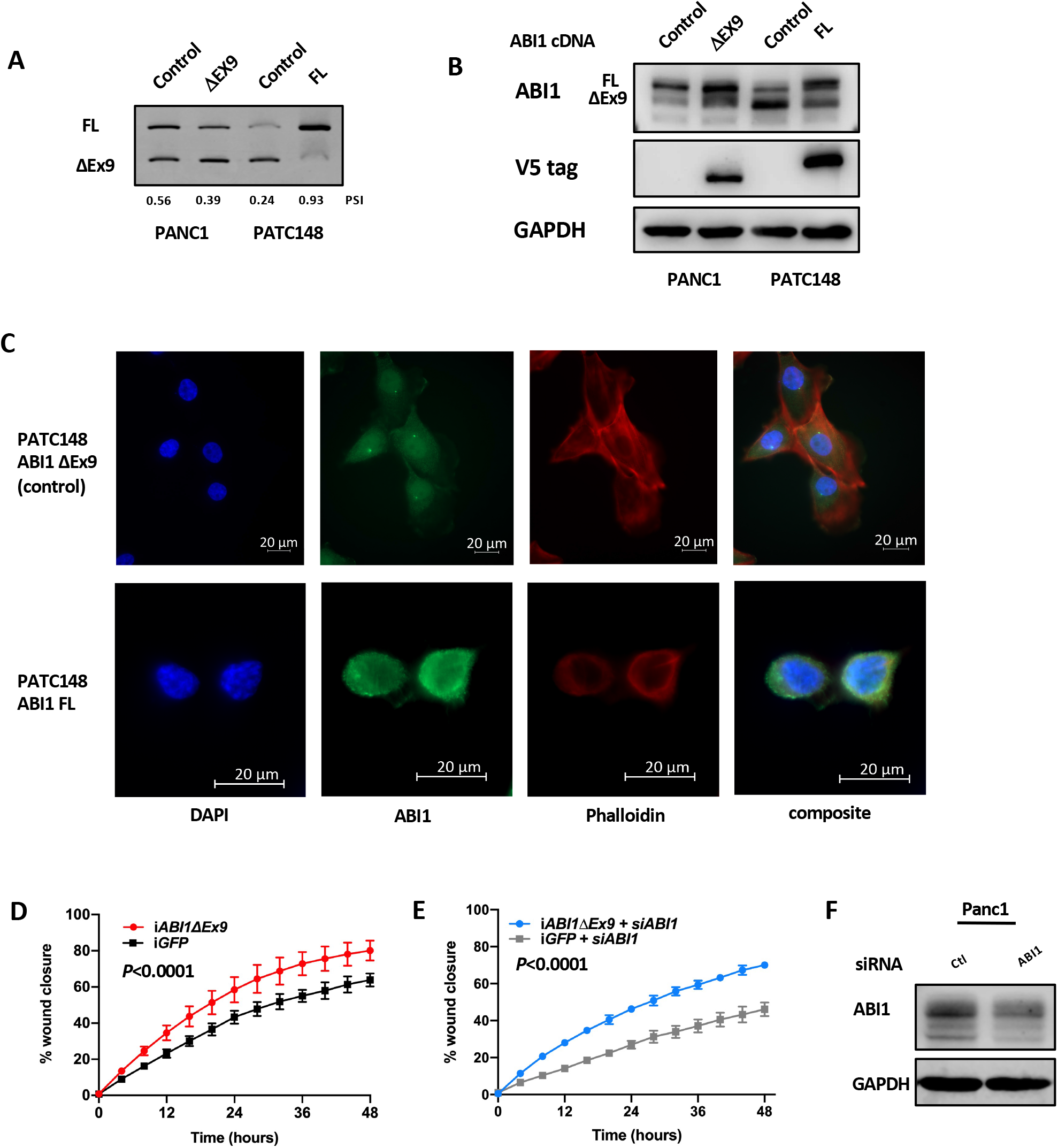
RBFOX2 loss promotes invadopodia formation through ABI1 splicing. Induced expression of ABI1 isoforms recapitulates splicing patterns observed with RBFOX2 repletion and depletion. Induced expression of ABI1ΔEx9 in RBFOX2-replete Panc1 cells reduces the PSI for ABI1, while induced expression of ABI1 FL in low RBFOX2-expressing PATC148 cells increases the ABI1 PSI to nearly exclusively ABI1 expression (**A**). Western blot analysis confirms the shift in ABI1 protein isoform expression. The V5 tag confirms the origin of the expressed isoforms is from the cDNA (**B**). PATC148 cells with the control vector show F-actin positive projections, with ABI1 distributed throughout the cytoplasm and in perinuclear regions (**C**, arrowhead). Cells with induced expression of ABI1 FL are contracted, with concentrated ABI1 and F-actin staining surrounding the perinuclear region. Induction of *ABI1*ΔEx9 in RBFOX2-replete Panc1 cells increases cell migration in a wound healing assay compared to induced control vector (**D**, *P*<0.0001, 2-factor ANOVA). Induced *ABI1*ΔEx9 rescues decreased cell migration observed in Panc1 cells with siRNA mediated *ABI1* knockdown. siRNA targeting the 3’ UTR of *ABI1* results in ∽50% knockdown of ABI1 protein in Panc1 cells (**F**).

### Splice-switching antisense oligonucleotides promote non-adherent growth and spheroid formation

Finally, to investigate whether ABI1ΔEx9 promotes other cellular phenotypes promoting pancreas cancer progression, we further investigated the function of the exon skipped isoform using splice switching antisense oligonucleotides (AONs) which target pre-mRNA transcripts containing *ABI1* exon 9 by binding the target exon and preventing processing of these transcripts by sterically inhibiting the splicing machinery (**Figure 9, panel A**) [14]. AONs were administered to Panc1 cells by free uptake from the media using an AON targeting ABI1 full length. Surprisingly, we observed that cells plated in the presence of the AON targeting *ABI1* did not form a uniform monolayer, instead forming dense clusters with low adherence (**Figure 9, panel B**). Trypan blue counting of these cells revealed 97-99% viability, suggesting that the AONs were not inducing cell toxicity. Cells treated with a non-targeting control AON at the same concentration formed the expected uniform monolayer. We confirmed the sustained enrichment of *ABI1ΔEx9* transcripts after 96 hours of AON free uptake by RT-PCR (**Figure 9, panel C**). Finally, because these cells failed to adhere uniformly to plastic culture plates, we asked whether the enrichment of ABI1 transcripts lacking exon 9 conferred more stem-like properties to the cells.

**Figure 9.**
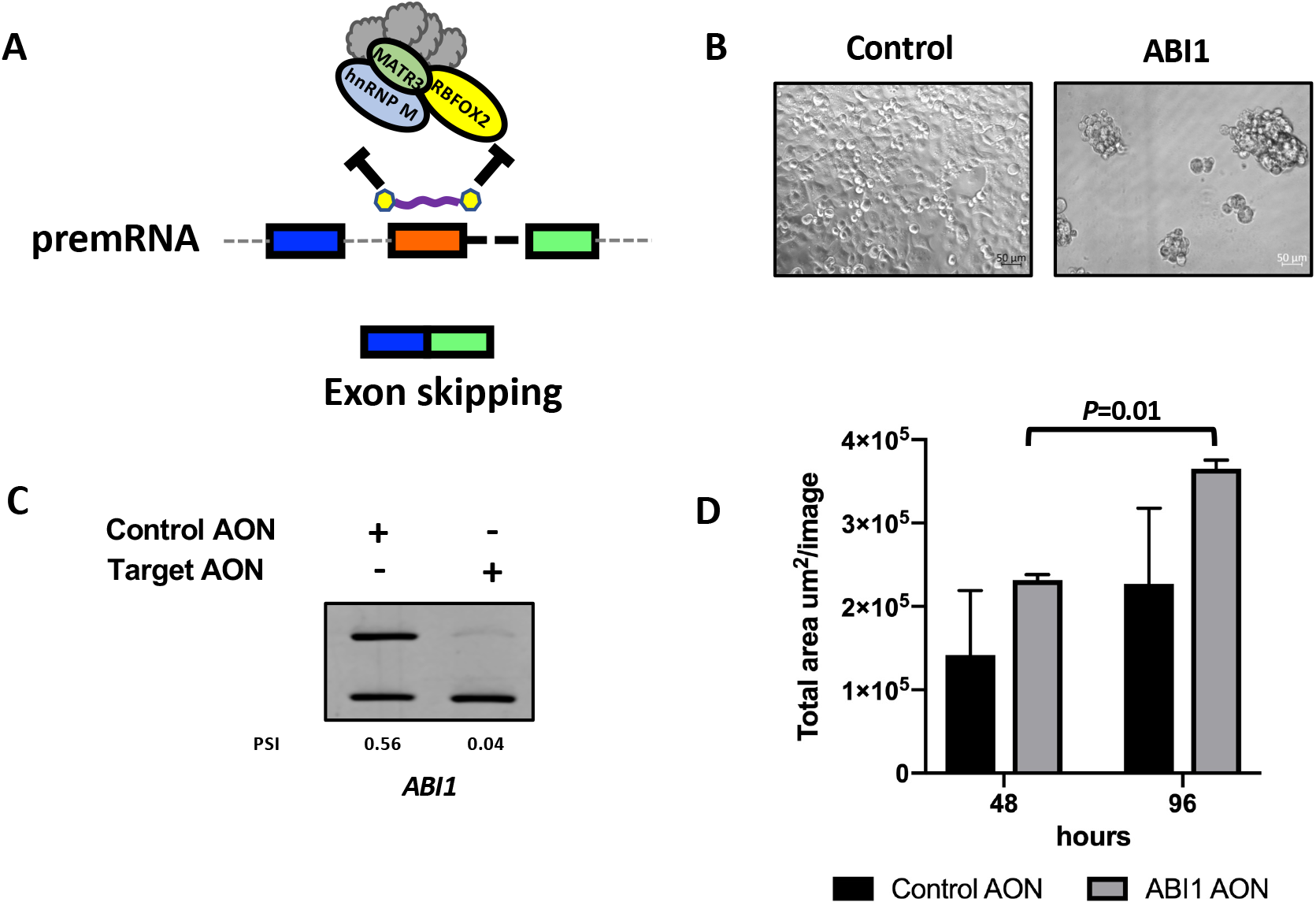
Induced ABI1ΔEx9 by splice-switching oligonucleotides promotes spheroid formation in Panc1 cells. Splice-switching antisense oligonucleotides (AONs) were designed to target pre-mRNA *ABI1* transcripts with exon 9 inclusion to prevent 5’ and 3’ splicing (**A**). Panc1 cells plated with 1 uM of a control AON formed an adherent monolayer in 2D culture on plastic. Panc1 cells plated with 1 uM AON targeting *ABI1* did not adhere to the plate and formed cell clumps (**B**). Cells were 90-95% viable after 48 hours. ABI1 exon 9 is excluded in Panc1 cells treated with the *ABI1* AON by free uptake (**C**, PSI 0.04) compared to cells treated with a control AON (PSI 0.56). Panc1 cells treated with AONs by free uptake were plated on 1:1 matrigel for spheroid assays. Induced and exclusive expression of the *ABI1*ΔEx9 transcript significantly increased the number and total area of spheres in Panc1 cells compared to control cells (**D**, *P*= 0.03, unpaired t-test).

To assess this, we used an Incucyte spheroid assay and plated 4000 cells treated with AONs for 24 hours on top of a 1:1 volume of Matrigel:serum-free media. Spheroid growth was tracked over a 5-day period with continual exposure to the AONs by free-uptake. We found a significant increase in the number and area of spheroids with the *ABI1* AON compared to control AON (**Figure 9, panel D**, *P*=0.03, unpaired t-test). Together, these data are the first to demonstrate that the *ABI1ΔEx9* isoform potentiates stem cell like properties to pancreatic cancer cells.

## DISCUSSION

Alternative splicing is an integral biological process promoting cellular plasticity, differentiation, and transformation. Recent pan-cancer analyses of splicing signatures suggest alternative splicing is a cancer hallmark and that cancers of epithelial origin exhibit conserved splicing events likely important for cancer progression [6, 7]. Mutation analysis of RNA binding proteins in cancer has elucidated oncogenic mutations in splicing regulators, including SF3B1 and U2F1, in some cancer types. In leukemias and lymphomas, these mutations are therapeutic liabilities [46]. However, splicing regulators are rarely mutated in pancreatic cancer. Our forward genetic screen identified a significant enrichment of RBPs perturbed through non-mutational events important in PDAC progression [22, 23]. Gene expression analysis of RNA binding proteins (RBPs) suggests they are differentially regulated at the transcriptional and post-transcriptional level, including SF3B1 [47] and RBPs MBNL1 and RBFOX2, which are regulated by alternative splicing [14, 36]. While RBFOX2 has been linked to EMT in breast cancer, in this study, we identified deregulation of RBFOX2 in pancreatic cancer through alternative splicing of the exon encoding the nuclear localization signal, resulting in a ratio of transcripts encoding nuclear and cytoplasmic isoforms of *RBFOX2* that varies across PDAC cell lines and in patient samples. This results in variable expression of nuclear RBFOX2 protein across patient samples, and we showed a significant association between reduced nuclear RBFOX2 and poorly differentiated pancreatic cancer. Poorly differentiated PDAC is associated with worse outcomes.

Using a series of isogenic cell lines replete and depleted for RBFOX2, we demonstrated that downregulation of RBFOX2 promotes cell migration, invasion and anoikis resistance in the absence of proliferative changes. These cellular phenotypes are reversed upon the expression of nuclear RBFOX2 in low expressing cell lines. *In vivo*, RBFOX2 loss promotes increased pancreatic tumor volume and metastatic incidence, particularly to the liver. Using expression arrays, we defined 45 differentially spliced exons controlled by RBFOX2 abundance, with an enrichment in events occurring in transcripts encoding proteins with known roles in cytoskeletal remodeling. These alternatively spliced transcripts encode a network of protein-protein interactions (PPI) surrounding RAC1, a known player in cell invasion [48-51]. Importantly, most of these splicing events lead to exon exclusion of conserved exons that are alternatively spliced across multiple cancers, including lung, breast and colon cancers [6-10]. Here, we showed for the first time that these splicing events also occur in pancreatic cancer, with a mean PSI below 0.5 for several exon splicing events, including *ABI1* and *DIAPH2*, suggesting that exon exclusion for RBFOX2 target exons is an integral part of pancreatic cancer progression. We found that exon inclusion for these targets correlated with RBFOX2 expression in PDAC patient samples, confirming that RBFOX2 plays a role in alternative splicing of these exons. Importantly, other splicing factors are likely to contribute to the complex regulation of exon splicing in pancreatic cancer. In fact, several of the exon splicing events we identified with RBFOX2 depletion were identified to be alternatively spliced by the RBP QKI in breast cancer cell lines [52]. *QKI* expression is low across most of the PDAC cell lines we assayed and may contribute to exon skipping of RBFOX2 target exons. Other splicing factors, including MBNL1, identified in our forward genetic screen [22, 23] may play a role in regulating alternative splicing in PDAC. Alternative splicing of *KIF13A*, an event we identified in our RBFOX2 depletion studies, was previously shown to be affected by MBNL1 abundance [12]. Exon inclusion of the RBFOX2 target exon in *ECT2* transcripts is low across PDAC cell lines, with RBFOX2 replete cell lines MiaPaCa2, Panc1 and 4039 exhibiting a PSI less than 0.5. *ECT2* encodes a small GTPase implicated as an oncogene in breast and other cancers [53-57] and RBFOX2-mediated exon skipping removes one of two exons encoding the N-terminal BRCT domain which negatively regulates ECT2 GTP binding [58]. RBFOX2 repletion experiments confirmed RBFOX2 has a role in controlling exon inclusion, but there is likely functional redundancy with other splicing factors regulating the function of ECT2 in cancer.

In this study, we show for the first time that RBFOX2 controls alternative splicing of *ABI1* and *DIAPH2* in pancreatic cancer. *DIAPH2* encodes a formin protein, and RBFOX2-mediated splicing removes an exon encoding part of the GTP binding domain of the protein, which may facilitate Rho-independent activation [31]. *ABI1* encodes the Ableson-like interacting protein 1 (ABI1), a protein that is part of the WAVE regulatory complex (WRC) that regulates actin remodeling through the interactions with the Arp2/3 complex to promote lamellopodia formation [59, 60]. We demonstrated that RBFOX2 depletion promoted lamellopodia formation in PDAC cells, redistributing F-actin to the cell periphery and increasing the number of F-actin foci. ABI1 protein relocalized to the cell periphery upon RBFOX2 depletion and overlapped with F-actin. Studies in breast cancer cells have shown reduced invadopodia formation and ECM degradation activity upon depletion of *ABI1* or *DIAPH2* [34]. However, the function of specific isoforms in cancer are understudied. Using a combination of inducible cDNAs and splice-switching antisense oligonucleotides (AONs), we uncovered a role for ABI1ΔEx9 in promoting pancreatic cancer progression by promoting cell migration and enhancing spheroid formation in Panc1 cells. These data suggest that the ABI1ΔEx9 isoform may confirm stem-cell like properties to these cells and warrants further investigation for its role in promoting pancreatic cancer progression and metastasis.

Alternative splicing is an important regulatory process in cancer. In this study, we elucidate the role of alternative splicing in promoting pancreatic cancer progression through deregulation of the splicing factor RBFOX2. While analyses to date have identified few splicing regulators with prognostic implications in PDAC, the processes regulated by alternatively spliced transcripts are integral to cancer progression. Alternative splicing signatures hold promise as biomarkers for cancer progression and metastasis and for potential therapeutic opportunities by targeting spliced transcripts directly to prevent translation or to identify protein isoforms with increased oncogenic potential conferred by alternative splicing that may be therapeutic targets [21]. Our studies lay the foundation for exciting avenues investigating the role of spliced isoforms in promoting cancer progression and metastatic potential in pancreatic and other cancers.

## Methods

### Cell culture

4039, Panc1, MiaPaCa2 and PL45 cell lines were obtained from ATCC and were propagated according to ATCC recommended conditions. 8902 cells were obtained from the DSMZ-German Collection of Microorganisms and Cell Cultures GmbH. MDA-PATC148 cells were obtained from MD Anderson Cancer Center and cultured in RPMI media supplemented with 2 uM glutamine and containing 100ug/mL primocin (Invivogen, Cat. no. ant-pm-1). Cell lines were routinely tested for mycoplasma and validated using STR profiling and mutational analysis.

### Vectors and lentivirus production

pGIPZ lentiviral vectors containing shRNAs directed against *RBFOX2* (Dharmacon) were purchased as lentiviral preparations (Cat. no. VGH5523-200225575, VGH5518-200275809, VGH5518-200230034). Additional shRNAs against *RBFOX2* were designed using splashRNA [61] and cloned into the lentiviral PRRL vector as described [62]. See Supplemental Table 4 for sequences. DDK-MYC tagged *RBFOX2* cDNAs were purchased from Origene in pCMV6-Entry vector (Cat. nos. RC206846 for RBFOX2-v1 and RC222907 for RBFOX2-v4). cDNAs were first subcloned into pLenti (Origene Cat. no. PS100092) maintaining the DDK-MYC tag and then into Pentr4FLAG (Addgene Cat. no. 17423) before cloning into the lentiviral DOX-inducible destination vector pLIX403 (Addgene Cat. no. 41395) using LR Clonase II. cDNAs for *ABI1* full-length (Cat. GC-M0337-CF-GS) and *ABI1*ΔEx9 (Cat. no. GC-Z8779-CF-GS) were purchased from Genecopoeia in pShuttle vectors without stop codons and subsequently recombined using LR Clonase II into pLIX403 with an in-frame 3’-end V5 tag. Lentiviral DNA constructs were packaged using ABMgood packaging mix (Applied Biological Materials Cat. no. LV053) transfected into 293 FT cell lines using jetPRIME transfection reagent (Polyplus, Cat. no. 89129-922). Lentivirus was concentrated by ultracentrifugation in clear ultracentrifuge tubes (Beckman Cat. no. NC9146666) spun in a Beckman Ultracentrifuge OPTIMA L 70K rotor (SW32 TI) at 23000 rpm for 2 hours at 4°C. Lentiviral titers were obtained using ABMgood qPCR Lentivirus Titer Kit (Applied Biological Materials, Cat. no. LV900). All vectors were confirmed by restriction digest and Sanger sequencing. Transductions were performed according to Dharmacon guidelines with 8 ug/mL polybrene and stably selected using puromycin.

### siRNA and AON transfections

siRNAs were purchased from SIGMA targeting *RBFOX2* (on-TARGET siRNA pool, Cat. no. M-020616-02-0005), *ABI1* (3’ UTR, Cat. no. M-017905-01-0005) or non-targeting control (Cat. no. D-001210-01-05). AONs were designed to target RNA sites accessible for binding as previously described [14, 63, 64]. Transfections were performed using Lipofectamine RNAiMax reagent (ThermoFisher Scientific).

### Cellular Assays

Proliferation assays were performed for adherent cells using Cell Titer Blue (Promega, Cat. no. G8080) and quantified using Promega Glomax Discover plate reader after four hours of incubation with the reagent at 37 degrees Celsius, 5% O_2._ For non-adherent live cell assays, cells were plated in 24-wells coated with a 40 mg/mL solution of Poly-HEMA (Poly 2-hydroxyethyl methacrylate; Sigma Cat. no. P3932) in complete media. Every 24 hours, media was collected, cells pelleted by centrifugation, and resuspended in 100 uL 0.25% Trypsin-EDTA (Invitrogen, Cat. no. SH30042.01). Cells were counted using a hemacytometer in duplicate per time point. Wound healing assays were performed using the Incucyte Live-Cell Analysis System (Incucyte Zoom, Sartorius). Cells were plated in 96-well plates (Essen Bioscience, Cat. no. 4379) overnight to form a confluent cell layer. 6 replicate wells were plated per cell line per assay. 16 hours post-plating, cells were treated with 10ug/mL mitomycin C (Cayman chemical, Cat. no. 11435-5) for 2.5 hours to inhibit cellular proliferation. A scratch was made using Incucyte® Woundmaker Tool (Cat. no. 4563) and cellular migration was assayed 72-96 hours.

Percent wound confluence was determined using Incycyte Zoom software. Where applicable, pLIX403 and PRRL constructs were induced using 250 ng/mL doxycycline for five days prior to plating cells for cellular assays and treatment continued throughout the experiment. Chemotatic invasion assays were performed using 4039 cells modified for RBFOX2 expression with the Incucyte Live-Cell Analysis System (Incucyte Zoom). Cells were serum-starved 16 hours prior to plating with 1% bovine serum albumin (VWR, Cat. no. 0332-100G) added for protein supplementation. Cells were treated with Cell Trace Far Red (Invitrogen, Cat. no. C34572) to detect nuclei according to the manufacturer’s instructions prior to plating cells in serum-free media with 1% BSA on top of Matrigel coated wells of Incucyte® Clearview 96-well plates (Corning Matrigel Growth Factor Reduced (GFR) Basement Membrane Matrix (Fisher Scientific, product no. CLS356231). 3D invasion into the matrix was monitored for 96 hours and percent invasion was calculated using the Integrated Incucyte® Chemotaxis Analysis Software Module (Sartorius). Spheroid assays were performed using Panc1 cells modified for RBFOX2 expression or target exon splicing modifications. 4E+03 cells were plated on 96-well plates (Corning, Cat. no. 3596) coated with 1:1 volume Matrigel:media prepared according to manufacturer’s recommended protocol for multispheroid assays. Spheroid growth in complete media was monitored for 5 days. The total area occupied by spheroids (total area um^2^/image) was calculated using the Incucyte® Spheroid Analysis Software module (Sartorius).

### Western blotting

Whole cell protein lysates from adherent or non-adherent cell cultures were isolated in RIPA buffer (100 mM Tris pH7.5, 300mM NaCl, 2mM EDTA, 2% NP-40, 2% Sodium deoxycholate and 0.2% SDS including protease inhibitors (Aprotinin (TOCRIS Bioscience Cat. no. 4139/10) Leupeptin (Fisher Scientific, Cat. no. 501685874) Sodium Orthovanadate (New England Biolabs, Cat. no. p0758s), Pepstatin (Cat. no. BP267110), Sodium Fluoride (Sigma, Cat. no. 201154-5G), AEBSF (Fisher Scientific, Cat. no. 40160000-2) and nuclease for cell lysis (Life Technologies, Cat. no. 88700) directly from tissue culture plates by scraping. Protein concentrations were determined using a BCA assay (Pierce, Catalog no 23227). Protein extracts were separated using SDS page gels with 4X Laemmli loading buffer and transferred onto nitrocellulose (BioRad Cat. No. 3500601) using a wet transfer apparatus and blocked using either LI-COR Intercept (TBS) Blocking Buffer (LI-COR, Cat. No. 927-60001) or 5% milk in TBST (20mM Tris pH 7.5,150mM NaCl with 0.1% Tween-20). Antibodies were diluted in the appropriate blocking buffer to visualize the following: RBFOX2 ((RBM9) Bethyl Labs, Cat. No. A300-864A); ABI1 (Cat. no. 39444), Vimentin (Cat. no. 5741S), CDH1(Cat. no. 3195S), ERK (Cat. no. 4695S), P-ERK (Cat. no. 4370S) AKT (Cat. no. 9272S), P-AKT-Ser473 (Cat. no. 9271S), JNK (Cat. no. 9252S), P-JNK (Cat. no. 9251S) and Rac1 (Cat. no. 4651) from Cell Signaling; GAPDH (Santa Cruz Cat. no. SC-69778); Beta-Actin (Cat. no A5441), Flag-M2 (Cat. no. F3165) and Histone H3 (Cat. no. 05-928) from Sigma. LI-COR Secondary antibodies for Infrared were IRDye 800cw anti-Rabbit (Cat. no. 926-32211) and IRDye 680RD anti-Mouse (Cat. no. 925-68070); HRP-conjugated anti-Rabbit (Vector Laboratories, Cat. no. PI-1000) and HRP-conjugated anti-Mouse (Jackson Immuno Research Labs, Cat. no. 115-035-003). HRP secondaries were developed using Pico chemiluminescent reagent (Pierce, Cat. no. PI34577). Gel images were acquired using LI-COR Odyssey and quantitated using ImageJ.

### Nuclear:cytoplasmic fractionation

PATC148 cells grown to 80% confluency were directly scraped off a 100 mm plate, pelleted and resuspended in 80 uL sucrose buffer with NP-40 (10mM TRIS pH 8.0, 320mM Sucrose, 3mM CaCl2, 4mM MgCl2, 0.1mM EDTA, 0.1%, NP40, 1mM DTT and protease inhibitors). The resuspension was transferred to a new 1.5 ml tube and centrifuged at 1500 RPM for 4 minutes in a swinging bucket rotor at 4°C. 60uL of the supernatant containing the cytoplasmic fraction was transferred to a new tube and kept on wet ice. The remaining supernatant was discarded and the cell pellet was washed twice with sucrose buffer without NP40. The nuclear pellet was resuspended in 80uL of complete RIPA buffer with protease inhibitors and nuclease. Exogenous Flag-tagged and endogenous RBFOX2 were probed in each cell fraction on an 8% SDS-PAGE gel. H3 histone and GAPDH were used as markers of the nuclear and cytoplasmic fractions, respectively.

### Rac1-GTP pulldown assays

Panc1 and 4039 cells grown to 80% confluency over 3 days were stimulated for 30 minutes with 30 ng/mL PDGF (Peprotech Cat. no. 100-00AB), then placed immediately on ice, washed once with PBS, then lysed and immediately scraped (Lysis Buffer: 50mM Tris pH 7.5, 10mM MgCl2, 0.5M NaCl, 2% NP-40 and protease inhibitors). The cell lysate was clarified by centrifugation at 10,000xg, 4°C for 1 minute, then transferred to a prechilled tube and quantitated by Bradford assay. 300 ug of protein in 600 uL of lysis buffer was used for the GTP pulldown with 20 ul of PAK beads (Sigma, Cat. no. 14-325) incubated for one hour. PAK beads were centrifugated at 5000xg at 4°C for 1 minute and washed once with 500 ul wash buffer (25mM Tris pH 7.5, 30mM MgCl_2_, 40mM NaCl). Washed beads were spun down and the bead pellet was resuspended in 20uL of 2x Laemmli Sample Buffer (BioRad, Cat. no. 161-0737) with 2-Mercaptoethanol (ICN Biomedicals, Cat. no. 806443). GTP-bound Rac1 and total Rac1 were visualized by western blot.

### Immunohistochemistry and immunocytochemistry

Analysis of RBFOX2 nuclear abundance was performed using immunohistochemistry on formalin-fixed, paraffin-embedded mouse tissue (pancreas and liver) and on pancreas tissue microarrays (MCC-TMA1, MCC-TMA2, MCC-TMA3) generated at Moffitt Cancer Center between 2006 and 2012 (RBFOX2 (RBM9) IHC00199, Bethyl Labs). Slides were scanned using the Aperio Digital Pathology ScanScope Slide Scanner (Leica Biosystems) at 20X magnification. Positive pixel counts for nuclear and cytoplasmic staining of RBFOX2 in tumor cells were determined using slide segmentation with Aperio ScanScope software. To visualize cellular morphologies and protein distribution for live cells grown in culture, cells were grown on acid-washed coverslips (Fisher, Cat. no. NC0706236) coated with bovine gelatin (Sigma, Cat. no. G1393), fixed with 4% paraformaldehyde and stained with Alexa Fluor 594-labelled phalloidin (ThermoFisher, Cat. no. A12381) according to methods previously described [65]. To quantify F-actin foci, images were captured at 63X using Zeiss Axio Imager upright microscope and analyzed using ImageJ with the ComDet plug-in (https://github.com/ekatrukha/ComDet). To visualize ABI1 isoform localization, cells were grown as described and incubated with phalloidin and ABI1 antibody (MBL International, Cat. no. D147-3) with Alexa-488 conjugated secondary (Jackson Immuno Research Labs, Cat. no. 115-545-146).

### Splicing analysis using arrays

Transcriptome profiling to define RBFOX2 splicing signatures from isogenic cell line pairs 4039, Panc1 and MiaPaCa2 with RBFOX2 knockdown and non-targeting control were generated using human Affymetrix Clariom D Arrays (Life Technologies, Cat. no. 902922) with the standard input kit run on the Applied Biosystems GeneChip 3000 instrument. Total RNA was extracted from whole cells or from pancreatic tumors using Qiagen RNeasy Mini kit DNase I treatment (Cat. No. 74106). Samples were prepared with the GeneChip WT Plus Reagent kit according to manufacturer’s guidelines. Differential exon usage analysis between RBFOX2 “high” (control) and RBFOX2 “low” (knockdown) was performed using Affymetrix Transcriptome Analysis Console (TACX) software. Analysis of cell lines and tumor samples was performed independently.

### Gene Expression and Splicing Analysis from Human RNA-seq data

Aligned data (BAM format) were downloaded for data sets EGAD00001004548 and EGAD00001003584. Within each data set, gene-level and exon-level expression count data were generated for each sample using the featureCounts command from the Subread software package [66] against GRCh38 v84 annotation and normalized to Transcripts Per Million (TPM). The Python script dexseq_prepare_annotation.py provided with R package DEXSeq [39] was used to translate Ensemble v84 GTF file to a GTF file with collapsed exon counting bin. The same GTF file was later used to calculate exon-level expression and PSI level. Percent spliced in index (PSI) was defined as the ratio between reads including or excluding an exon, indicating how frequently the exons of interest are spliced into transcripts. It was calculated following a previous published protocol [38]. For a given exon, PSI=1 means this exon is 100% spliced in; while PSI=0 means this exon is 100% spliced out. Pearson and Spearman correlation between RBFOX2 expression (TPM) and target exon PSI were calculated using R 4.0.3. Permutation test was performed to compare the average absolute correlation of the exons of interests (the true) to the average absolute correlation of randomly selected same number of exons (repeated 10,000 times). Empirical p-value was calculated as probability of (random average correlation > true average correlation). Splicing analysis from TCGA PAAD RNA-seq data was interrogated using SpliceSeq [37]. For data set E-MTAB-6830, FASTQ data were downloaded from the NCBI SRA, and aligned to the Human GRCh38 reference genome using STAR (version 2.7.3a) [67]. Gene-level count data were generated using featureCounts [66] based on GRCh38 v99 annotation. For both data sets, count data were converted to log_2_ counts per million, and processed within the R computing environment with the Voom normalisation procedure from the limma software package [68]. For EGAD00001004548 and E-MTAB-6830 datasets, Moffitt subtype information was generated per-sample using the following procedure: data for the 28 genes used to define the Moffitt subtypes were extracted from each data set, and consensus clustering was then used to separate the samples into two distinct clusters (Basal-like and Classical). Cluster identify was confirmed by checking survival associations (in each case the Basal-like subtype was associated with worse prognosis, as per the original paper) [31]. Additional clinical information was obtained for the EGAD00001004548 data set by matching sample IDs to those available from the ICGC PACA-CA project (matches were found for 197 samples).

### Real-time and qRT-PCR

RNA was extracted from whole cells using EZNA total RNA kit (R6834-02, Omega Bio-Tek). For gene expression analysis, cDNAs were made using 1ug of extracted RNA using qScript™ cDNA Synthesis Kit (Cat. no. 95047-100). qPCR was carried out using PerfeCTa® SYBR® Green SuperMix Reaction Mix (Cat. no. 95055-500) in real time PCR system Quant Studio 7 PRO (Applied Biosystems). For RT-PCR analysis of splicing, 1 ug RNA was used to prepare cDNA with the Vilo cDNA kit (Invitrogen Cat. No. 11755050). PCR amplification was performed with Roche Taq (Cat. No. 11146173001) and PCR products were visualized on 12% acrylamide gels stained with 5 ug/mL ethidium bromide. Gel images were captured using Licor Odyssey FC and PCR bands quantified using Image Studio Lite software. PSI calculations were performed as previously described [9, 12]. Primers sequences are provided in Supplemental Table 4.

### Mice

Human 4039 (5×10^4), Panc1 (1×10^5) and PATC148 (1×10^5) cells were orthotopically introduced into the pancreas of male and female NSG recipient mice (JAX 005557) according to methods described previously [69]. Isogenic cell lines (knockdown vs. control) were matched for cell passage, implanted into littermates and aged concurrently. Mice were aged for 45 days. Primary tumor volumes, metastatic incidence and lesion sizes were measured at necropsy. Pancreas tumors were formalin fixed and embedded with the spleen attached for orientation. A portion of the tumor was snap frozen for molecular analysis. Metastatic lesions greater than 2 mm were formalin-fixed for histology. All procedures were conducted under approved IACUC protocols and following AALAC guidelines.

### Statistical Analysis

Statistical analysis of cell-based assays and *in vivo* tumor growth was carried out using GraphPad PRISM software. Individual statistical tests are referenced within figure legends. Oncoprints were generated from published datasets from the Sleeping Beauty Cancer Driver Database (SBCDDB, [23]) using the cBioPortal Oncoprinter tool.

## Author contributions

K.M.M. designed and supervised the study; M.M. and K.M.M. performed gene knockdown studies, overexpression studies and immunocytochemistry; M.M., K.M. and R.M. performed splicing analysis; M.M. and K.M. carried out the cloning of all constructs; K.M.M., and M.R. performed proliferation and migration assays; M.M. and M.R. performed western blot analysis; K.M.M. performed orthotopic surgeries; J.B.F provided cell lines and patient samples; B.C. provided pathology expertise; E.G. provided splicing expertise and AON design; X.Y., J.Y.N. and M.A.B performed bioinformatic analyses of splicing in human pancreatic cancer RNA-seq datasets. K.M.M. wrote the manuscript with input from M.M, M.A.B. and E.G. All authors read and approved the final manuscript.

## Acknowledgements

We are indebted to the expertise of Sean Yoder and the Moffitt Cancer Center Molecular Genomics Core, Eric Welsh and the Moffitt Cancer Center Biostatistics and Bioinformatics Core, Joe Johnson and the Moffitt Cancer Center Microscopy Core, and Noel Clark and the Moffitt Cancer Center Tissue Core supported by CCSG P30-CA076292. Shengyu Yang (Penn State College of Medicine) provided protocols and advice for invadopodia immunocytochemistry. Antje Schaefer and Channing Der (UNC Chapel Hill) provided protocols and advice for Rac1-GTP pulldown assays. We thank Margi Baldwin, Haley Baldwin and the University of South Florida Comparative Medicine facility for assistance with *in vivo* studies. We are grateful for the help of Michael Mann for analysis of Sleeping Beauty datasets. Finally, we thank Marco Napoli and Elsa Flores, Moffitt Cancer Center, for their generous support and expertise with Incucyte assays.

