## Supplemental Figures for "RBFOX2 deregulation promotes pancreatic cancer progression and metastasis through alternative splicing"

### SUPPLEMENTAL FIGURE LEGENDS

**Supplemental Figure 1. Identification of *Rbfox2* as a progression driver in a Sleeping Beauty mouse model of pancreatic cancer.** Progression driver genes identified in an *in vivo* forward genetic screen using *Sleeping Beauty* (SB) insertional mutagenesis in a GEMM model of pancreatic cancer are statistically enriched for regulation of mRNA splicing via the spliceosome. An oncoprint of the SB transposon insertions in regulators of mRNA splicing identified *Mbnl2*, *Mbnl1* and *Rbfox2* as the most frequently mutated genes (**A**). Expression analysis of the top three splicing factors in tumor vs. normal pancreas from GEO dataset GSE28735 showed a significant decreased in *MBNL2* expression in tumors (**B,** FDR adj. *P*=0.0321) and a significant increase in *RBFOX2* expression in tumors (**C,** FDR adj. *P*=0.006) compared to normal pancreas, while *MBNL1* showed no significant difference in gene expression (**D**, FDR adj. *P*=0.2299). Quantitative PCR (qPCR) analysis of *MBNL2* (**E**) and *MBNL1* (**F**) lines shows variable expression in human pancreatic cancer cell lines, with the highest levels in Panc1 and 8902 cells.

**Supplemental Figure 2. Cellular signaling in RBFOX2-depleted cells**. Panc1 and MiaPaCa2 cells grown under non-adherent conditions for 48 hours exhibit activation of JNK and AKT, two cell survival markers. JNK phosphorylation is slightly elevated in Panc1 and MiaPaCa2 cells depleted for RBFOX2 (**A**) compared to cells replete for RBFOX2 and is quantified over three replicate experiments in (**B**). AKT Ser473 phosphorylation is slightly elevated in Panc1 cells depleted for RBFOX2 compared to cells replete for RBFOX2 while phospho-AKT levels are unchanged in MiaPaCa2 cells (**C**) and quantified in (**D**). Under adherent growth conditions, phospho-ERK1 (upper band) is depleted in Panc1 cells with RBFOX2 and is not expressed by MiaPaCa2 cells. Levels of total ERK or phospho-ERK2 (P-ERK, lower band) are unchanged across all isogenic pairs (**E**).

**Supplemental Figure 3. Inducible *RBFOX2* knockdown recapitulates cellular phenotypes.** A DOX-inducible shRNA targeting *RBFOX2* decreased RBFOX2 protein expression (**A**) in the absence of changes in mesenchymal-like properties of the cells as assessed by expression of CDH1 (E-cadherin) and vimentin (VIM). Quantification of RBFOX2 protein reduction is shown for cells with inducible *RBFOX2* knockdown compared to inducible non-targeting shRNA control (**B**). Inducible *RBFOX2* depletion does not change cellular growth under adherent conditions (**C**). MiaPaCa2 cells with inducible *RBFOX2* knockdown demonstrate significantly increased survival under non-adherent conditions compared to control cells (**D**, 2-factor ANOVA *P*=0.0015). 4039 cells show significantly increased cellular migration with inducible RBFOX2 depletion (**C**, 2-factor ANOVA *P*<0.001). Cells were treated with doxycycline for 5 days prior to analysis.

**Supplemental Figure 4. Inducible expression of cytoplasmic RBFOX2 v4 isoform does not alter cellular phenotypes.** Induced over-expression of the RBFOX2 v4 isoform in PATC148 and 8902 cell lines is demonstrated by western blot analysis of total RBFOX2 and the Flag tag (**A**). Expression of CDH1 and vimentin are unchanged upon expression of RBFOX2 v4. Quantification of RBFOX2 v1 and v4 isoforms in PATC148 and 8902 cells compared to GFP control shows similar expression levels of the different isoforms in each isogenic series (**B**). Nuclear:cytoplasmic separation of PATC148 cells with an induced GFP control vector ("GFP") confirms that RBFOX2 expression is low in both the cytoplasmic ("C") and nuclear ("N") fractions (**C**). Induced over-expression of RBFOX2 v1 protein is detected in both the cytoplasm ("C") and nucleus ("N"). The predominance of the Flag tag in the nuclear fraction confirms the v1 isoform resides in the nucleus. Induced overexpression of the v4 isoform demonstrates a lower abundance of total RBFOX2 protein and exclusive localization in the cytoplasmic fraction, confirmed by the Flag tag. Western blot detection of GAPDH only in the cytoplasmic ("C") fraction and Histone H3 only in the nuclear ("N") fraction confirms clean nuclear:cytoplasmic fractionation. Expression of RBFOX2 v4 does not change the proliferative capacity of the PATC148 and 8902 cells (**D**). Expression of RBFOX2 v4 does not change the migratory capacity of PATC18 cells in a wound healing assay (**E**).

**Supplemental Figure 5. RBFOX2 depletion in Panc1 cells significantly increases PDAC progression.** Constitutive RBFOX2 knockdown in Panc1 cells significantly increased pancreas tumor volumes in an orthotopic model in NSG mice compared to animals injected with Panc1 cells with a non-targeting shRNA (**A**, unpaired t-test, *P*=0.011). RBFOX2 depletion also increased the incidence of macro metastasis to the liver (LV), mesentery (MES), spleen (SP) and kidney (KID) compared to Panc1 cells replete for RBFOX2, represented as the percentage of animals with at least one focus in the target organ measured as greater than 1 mm at necropsy (**B**). The volume of metastatic lesions ($mm^3$) collected at necropsy is graphed on a per-animal basis (**D**). Only one mouse from the control cohort exhibited liver lesions. H&E staining of a pancreas tumor from Panc1 cells expressing the non-targeting shRNA (**C**) shows tumor with surrounding normal pancreas (lower left corner). H&E staining of a pancreas tumor (**E**) and liver metastasis (**F**) from Panc1 cells with shRNA mediated *RBFOX2* knockdown. Analysis of RBFOX2 expression by immunocytochemistry demonstrates robust RBFOX2 expression in tumors from replete cells (**G**) and decreased signal in RBFOX2 depleted pancreas tumor (**H**) and resulting liver metastasis (**I**). Surrounding normal liver expresses low RBFOX2 protein.

**Supplemental Figure 6. RBFOX2 v1 isoform repletion in PATC148 cells reduces PDAC metastasis.** Induced expression of RBFOX2 v1 in tumors established using PATC148 cells does not change end-stage pancreatic tumor volumes compared to tumors with induced expression of a GFP control (**A**). The incidence of macro metastases is not observed in mice with induced RBFOX2 v1 in mesentery (MES), diaphragm (DIAPH), kidney (KID) and bodywall (**B**). H&E staining of pancreas tumors from RBFOX2 low (**C**) and RBFOX2 high (**D**) tumors shows little normal pancreas. RBFOX2 immunohistochemistry in RBFOX2-low (**E**) and RBFOX2 replete showed heterogeneous expression with similar RBFOX2 intensity.

**Supplemental Figure 7. RBFOX2 regulates alternative splicing of transcripts encoded by mouse orthologs identified from Sleeping Beauty PDAC mouse model.** Eleven exon splicing events controlled by RBFOX2 occur in transcripts encoded by statistically defined cancer genes in a Sleeping Beauty mouse model of PDAC. An oncoprint highlights the incidence of SB transposon insertions in these mouse orthologs, with "hits" depicted by blue bars and the percentage of tumors with insertions in a population of 172 tumors highlighted on the left-hand side.

**Supplemental Figure 8. RBFOX2 deregulation promotes alternative splicing of RBFOX2 target exons.** Real-time PCR (RT-PCR) validation of RBFOX2 target exon usage in PDAC cells with inducible knockdown of RBFOX2 using sh1443 confirms exon skipping of RBFOX2 target exons in *ABI1*, *DIAPH1*, *DIAPH2* and *ECT2* compared to cells with an inducible nontargeting shRNA (Ctl). Percent spliced-in (PSI) values are calculated for RBFOX2 target exons in each isogenic pair (**A**). The percent abundance of RBFOX2 protein upon induced RBFOX2 knockdown is shown in panel (**B**). Induced RBFOX2 v4 isoform expression in PATC148 and 8902 cells does not change the PSI of spliced target exons in *ABI1*, *DIAPH1*, *DIAPH2* and *ECT2* transcripts compared to control cells (**C**). The percent abundance of RBFOX2 protein upon induced RBFOX2 knockdown in 8902 cells (**D**). PSI values for RBFOX2 target exons decrease in 8902 cells with constitutive (sh145) or inducible (sh1443) RBFOX2 knockdown in 8902 cells compared to controls (**E**). Expanded analysis of RBFOX2 target exon splicing in PDAC by using either the constitutive shRNA targeting RBFOX2 (sh145) and the constitutive non-targeting shRNA control (Ctl) or the inducible sh1443 targeting RBFOX2 and the matched inducible non-targeting control (Ctl) shows parity for spicing shifts based on PSI values for each isogenic cell line pair (**F**). The degree of exon skipping with RBFOX2 knockdown corresponds with the increased abundance of the lower PCR product and reduced PSI value and is dependent on the degree of RBFOX2 knockdown. The majority of RBFOX2 target exons are skipped in the absence of RBFOX2, while for select transcripts *ADD3*, *EXOC1* and *MAP3K7*, RBFOX2 target exons are included upon reduced RBFOX2 expression.

**Supplemental Figure 9. Correlation of RNA abundance with exon inclusion for RBFOX2 target exons in human PDAC patient samples.** A histogram depicts the permutation test to compare the average absolute correlation between the PSI for spliced exons and *RBFOX2* expression with the average absolute correlation of randomly selected exons and *RBFOX2* expression in resected human PDAC patients (**A**) (mean Pearson correlation coefficient 0.1981, $P$=0.0009). The population correlation between the RBFOX2 target exon PSIs and RBFOX2 expression in pancreas liver metastases for *DIAPH1* exon 3 (**B**), ABI1 exon 9 (**C**) and *DIAPH2* exon 2 (**D**).

**Supplemental Figure 10. siRNA mediated RFOX2 knockdown to define ABI1 localization.** siRNA targeting *RBFOX2* in Panc1 and MiaPaCa2 cells achieved significant RBFOX2 protein reduction (**A**). si*RBFOX2* promoted a decreased PSI for ABI1 exon 9 compared to a nontargeting siRNA in both Panc1 and MiaPaCa2 cells (**B**). ABI1 protein isoforms in si*RBFOX2* treated cells shift towards ABI1ΔEx9 in Panc1 cells and a reduction in ABI1 full length in MiaPaCa2 cells (**C**). Immunocytochemistry for ABI1 in Panc1 cells shows increased expression with si*RBFOX2* compared to control and an increased signal at the cell periphery, coincident with F-actin (Phalloidin). Cells were grown on gelatin substrates.

### Supplemental Figure 1

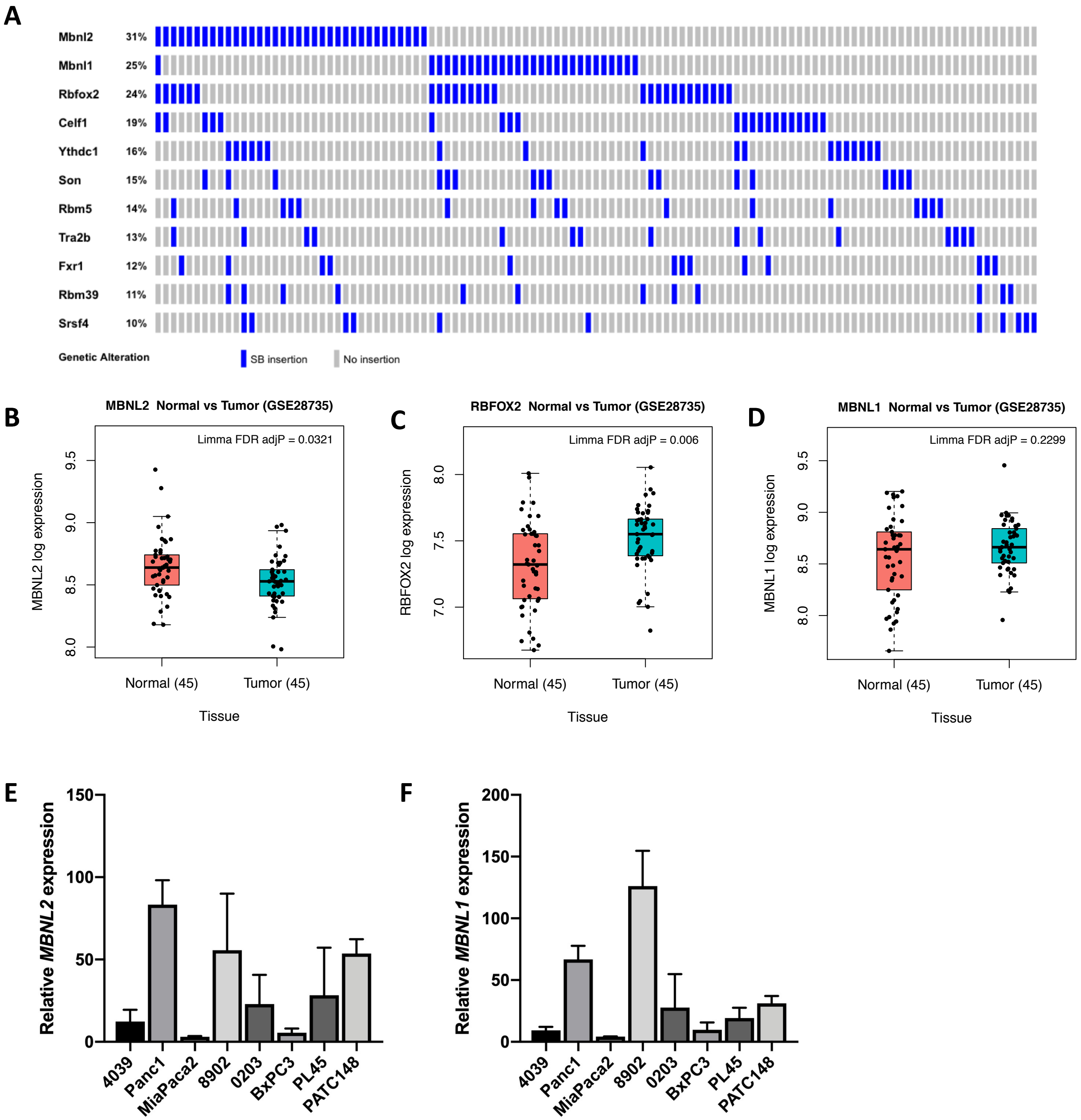

Supplemental Figure 2

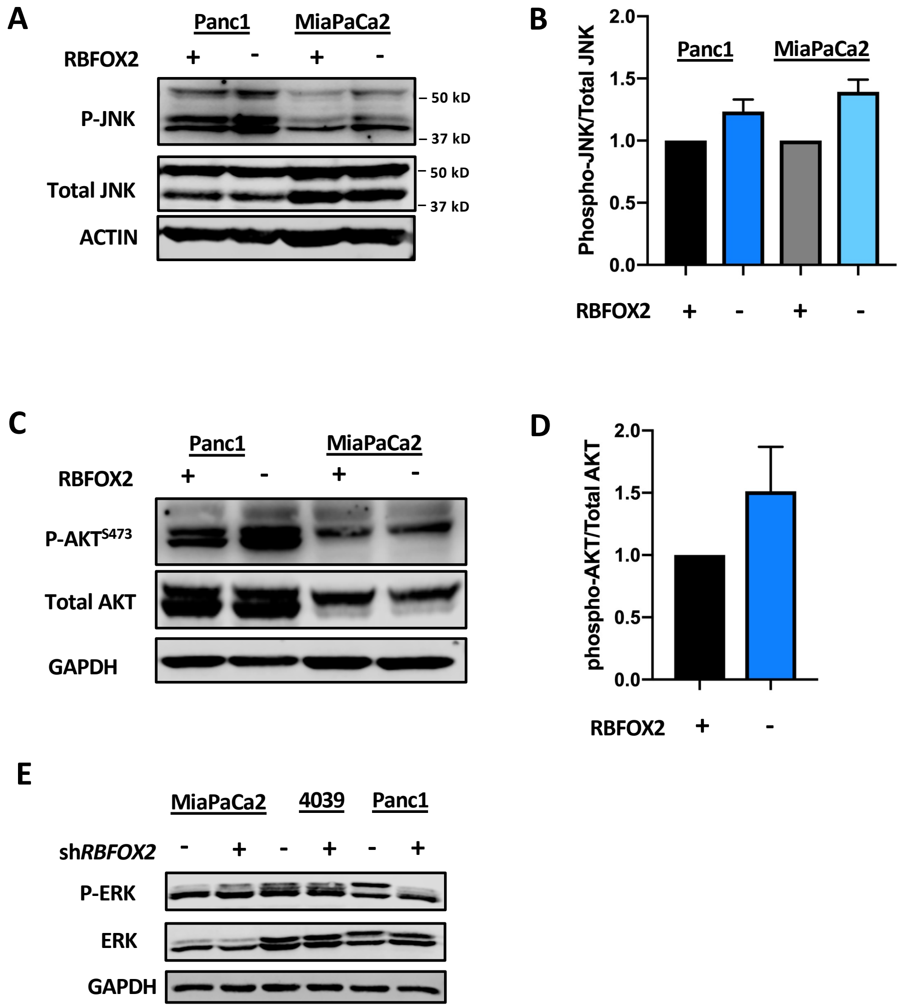

**Supplemental Figure 3**

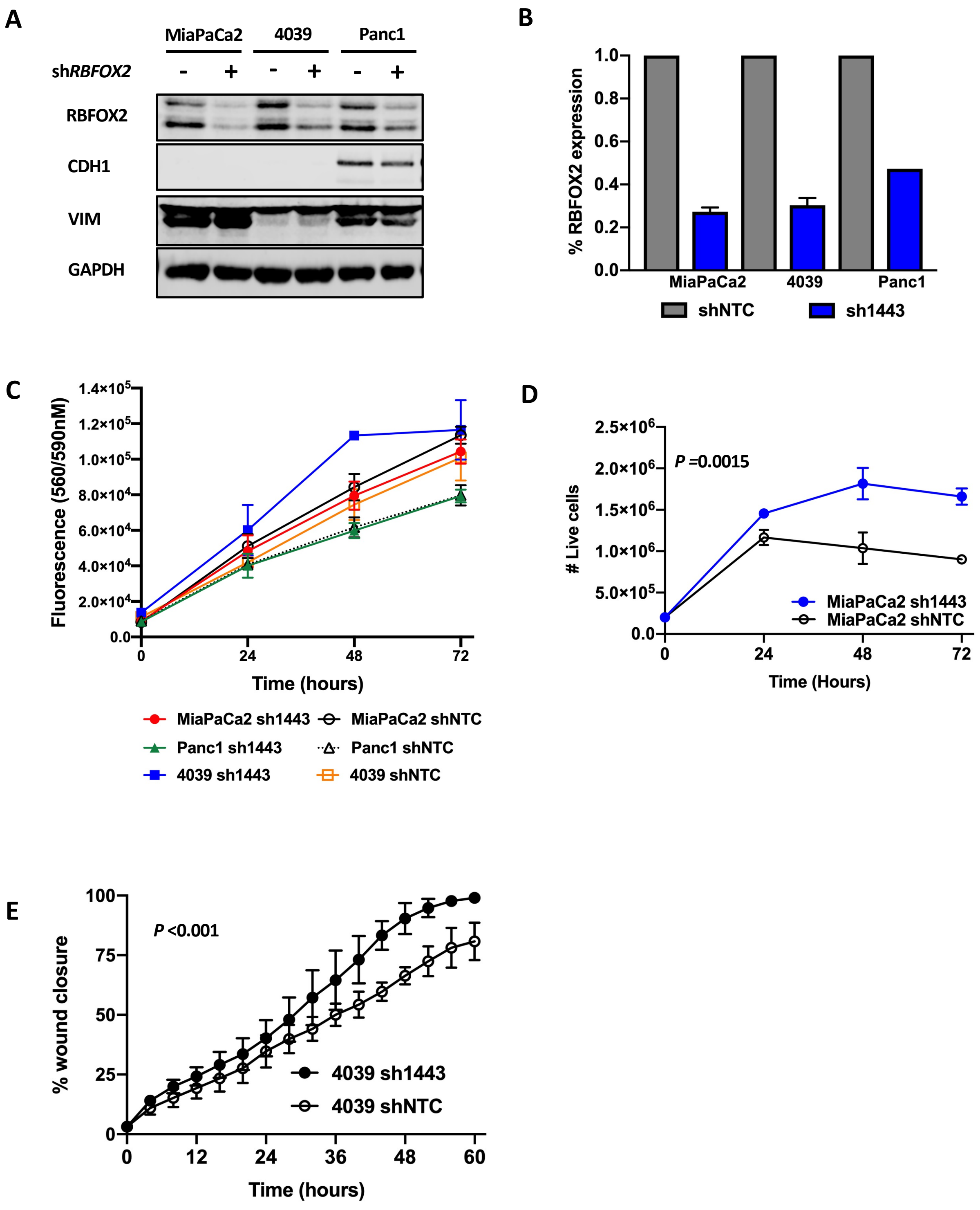

### Supplemental Figure 4

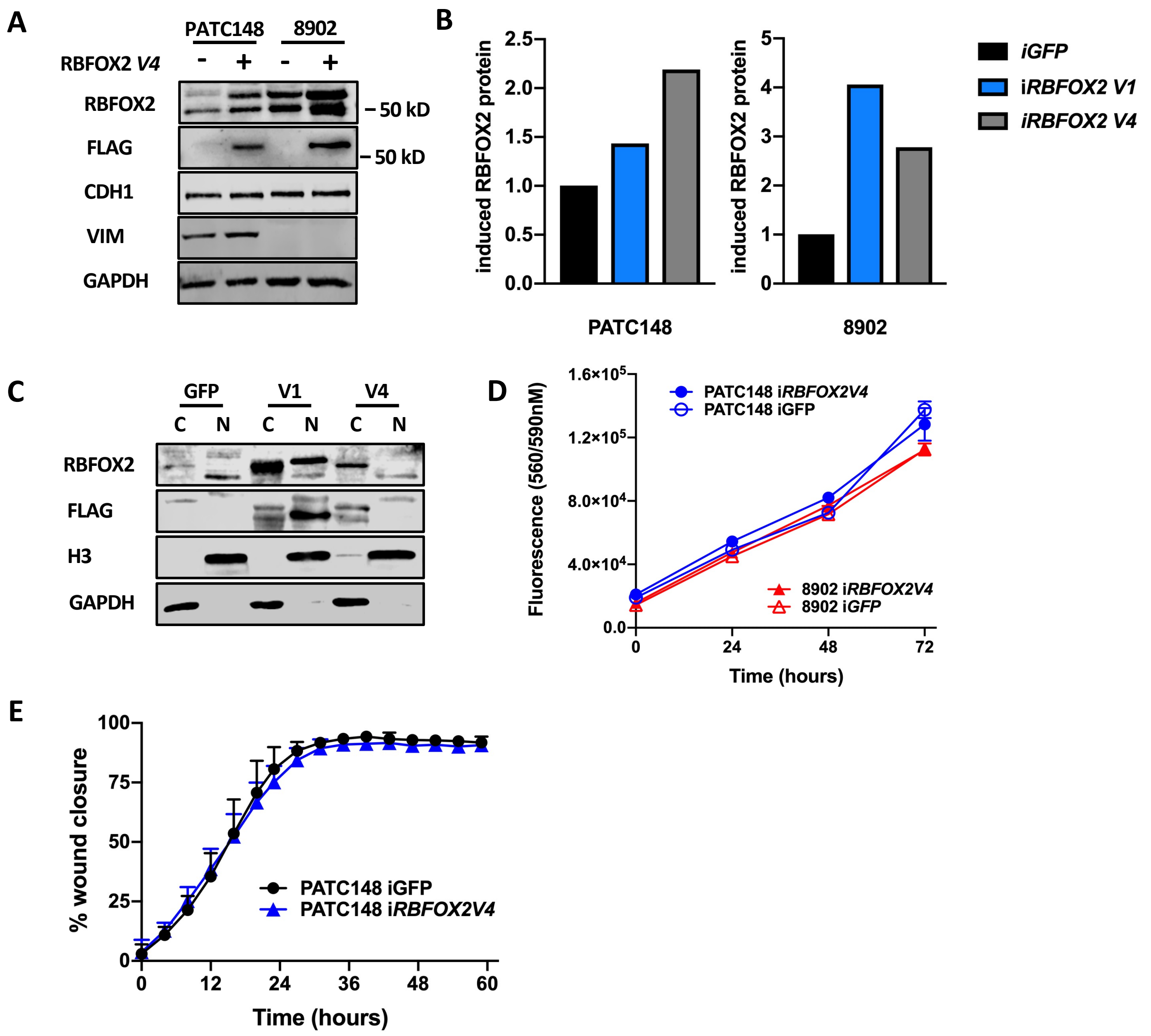

### Supplemental Figure 5

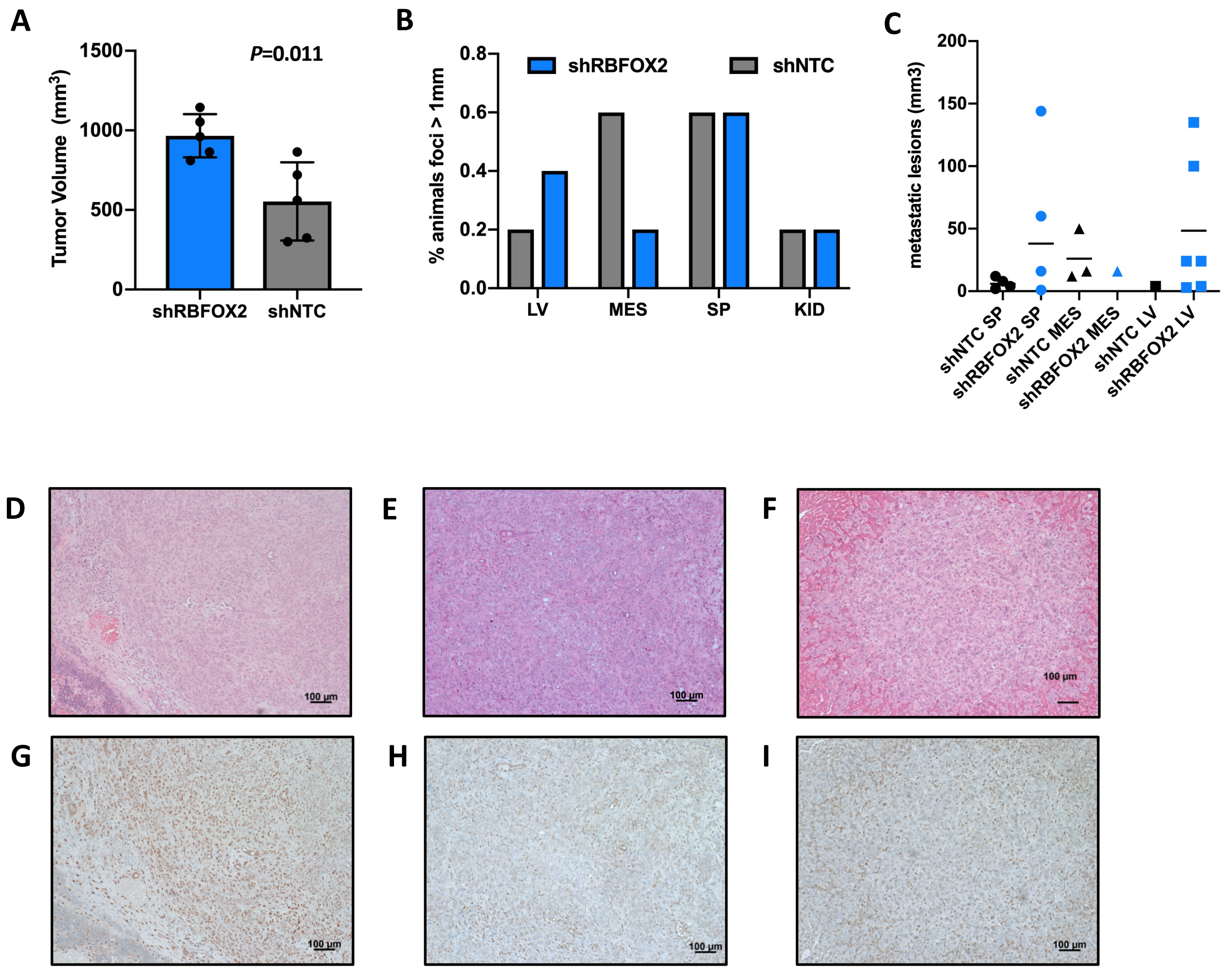

**Supplemental Figure 6**

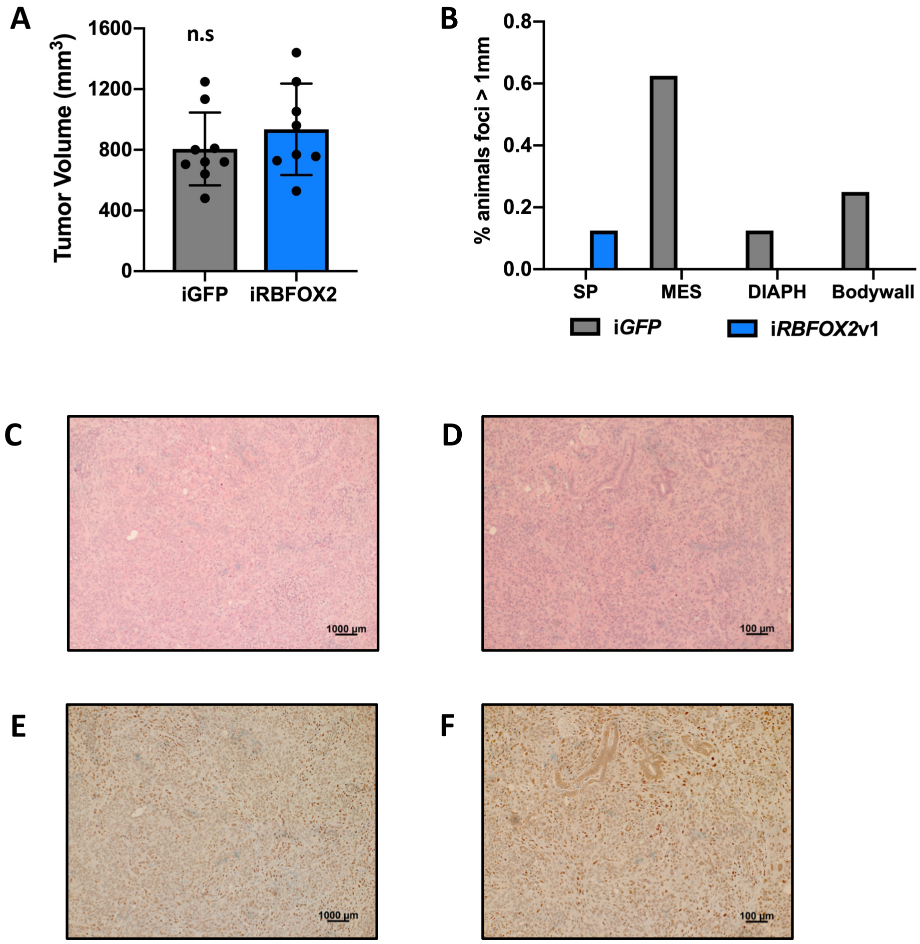

**Supplemental Figure 7**

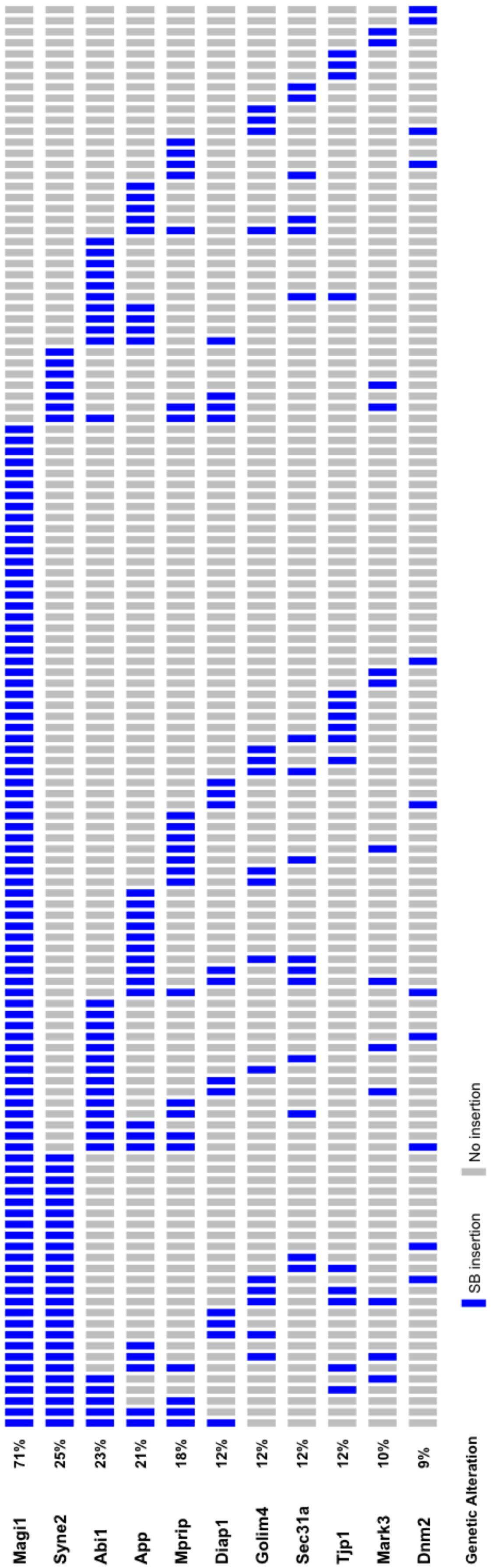

### Supplemental Figure 8

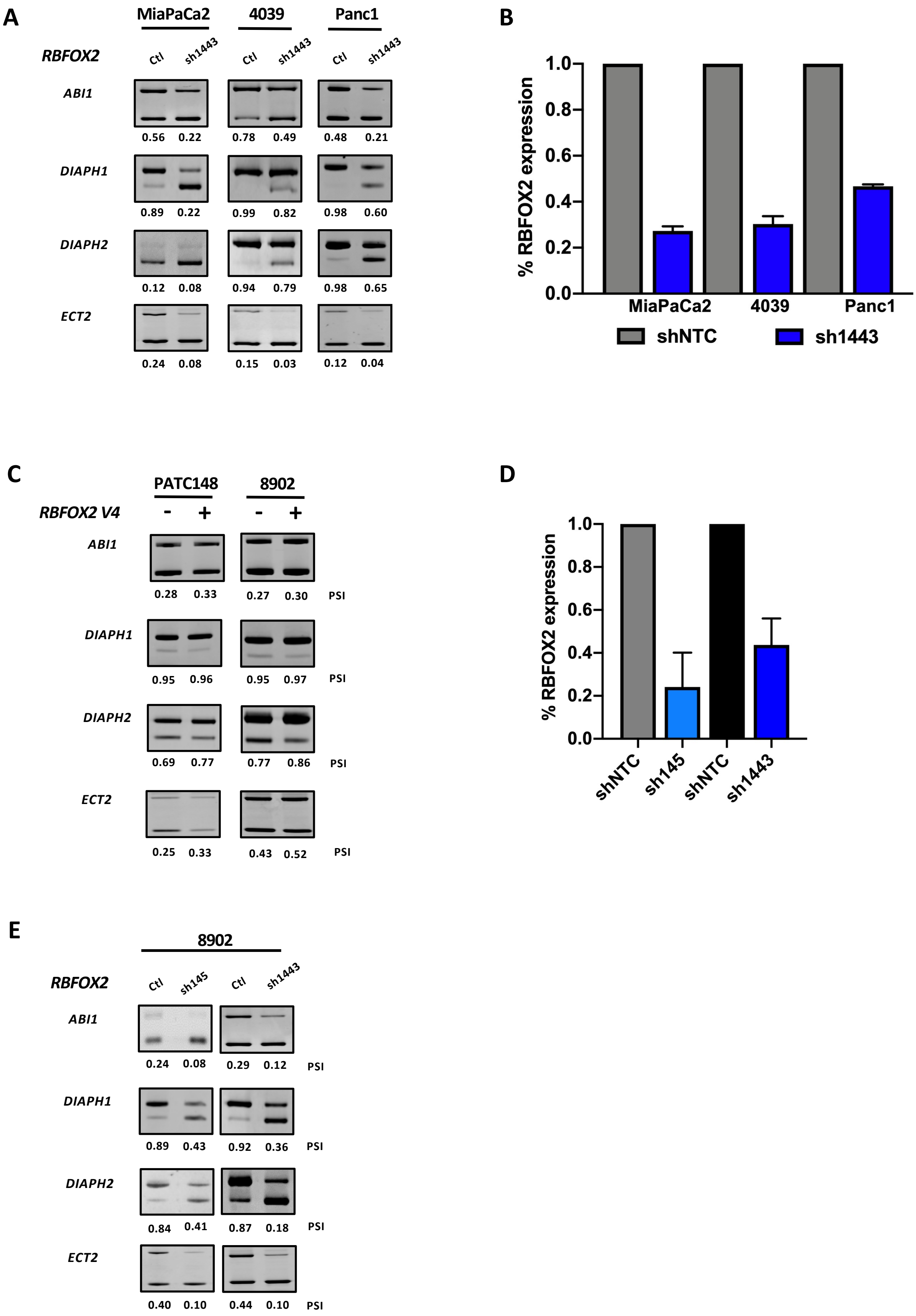

F

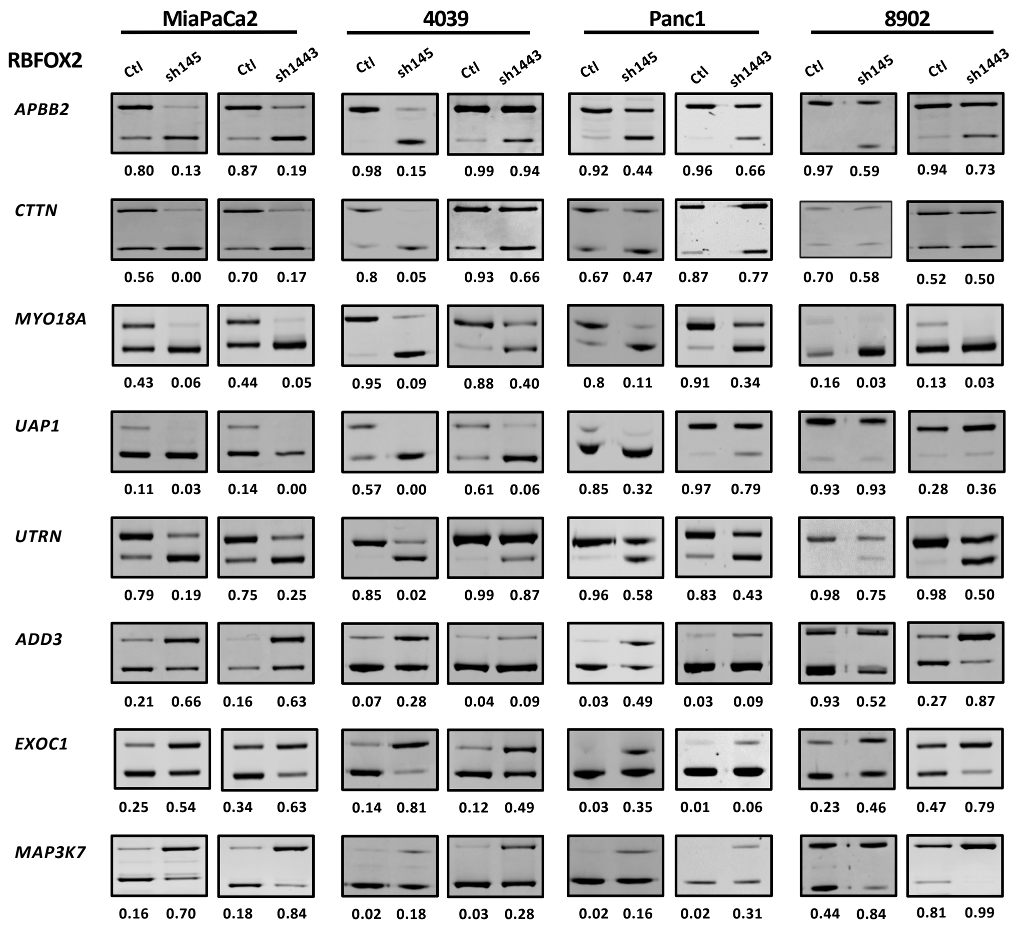

MiaPaCa2
4039
Panc1
8902
RBFOX2
Ctl
sh145
Ctl
sh1443
APBB2
0.80 0.13 0.87 0.19 0.98 0.15 0.99 0.94 0.92 0.44 0.96 0.66 0.97 0.59 0.94 0.73
CTTN
0.56 0.00 0.70 0.17 0.8 0.05 0.93 0.66 0.67 0.47 0.87 0.77 0.70 0.58 0.52 0.50
MYO18A
0.43 0.06 0.44 0.05 0.95 0.09 0.88 0.40 0.8 0.11 0.91 0.34 0.16 0.03 0.13 0.03
UAP1
0.11 0.03 0.14 0.00 0.57 0.00 0.61 0.06 0.85 0.32 0.97 0.79 0.93 0.93 0.28 0.36
UTRN
0.79 0.19 0.75 0.25 0.85 0.02 0.99 0.87 0.96 0.58 0.83 0.43 0.98 0.75 0.98 0.50
ADD3
0.21 0.66 0.16 0.63 0.07 0.28 0.04 0.09 0.03 0.49 0.03 0.09 0.93 0.52 0.27 0.87
EXOC1
0.25 0.54 0.34 0.63 0.14 0.81 0.12 0.49 0.03 0.35 0.01 0.06 0.23 0.46 0.47 0.79
MAP3K7
0.16 0.70 0.18 0.84 0.02 0.18 0.03 0.28 0.02 0.16 0.02 0.31 0.44 0.84 0.81 0.99

**Supplemental Figure 9**

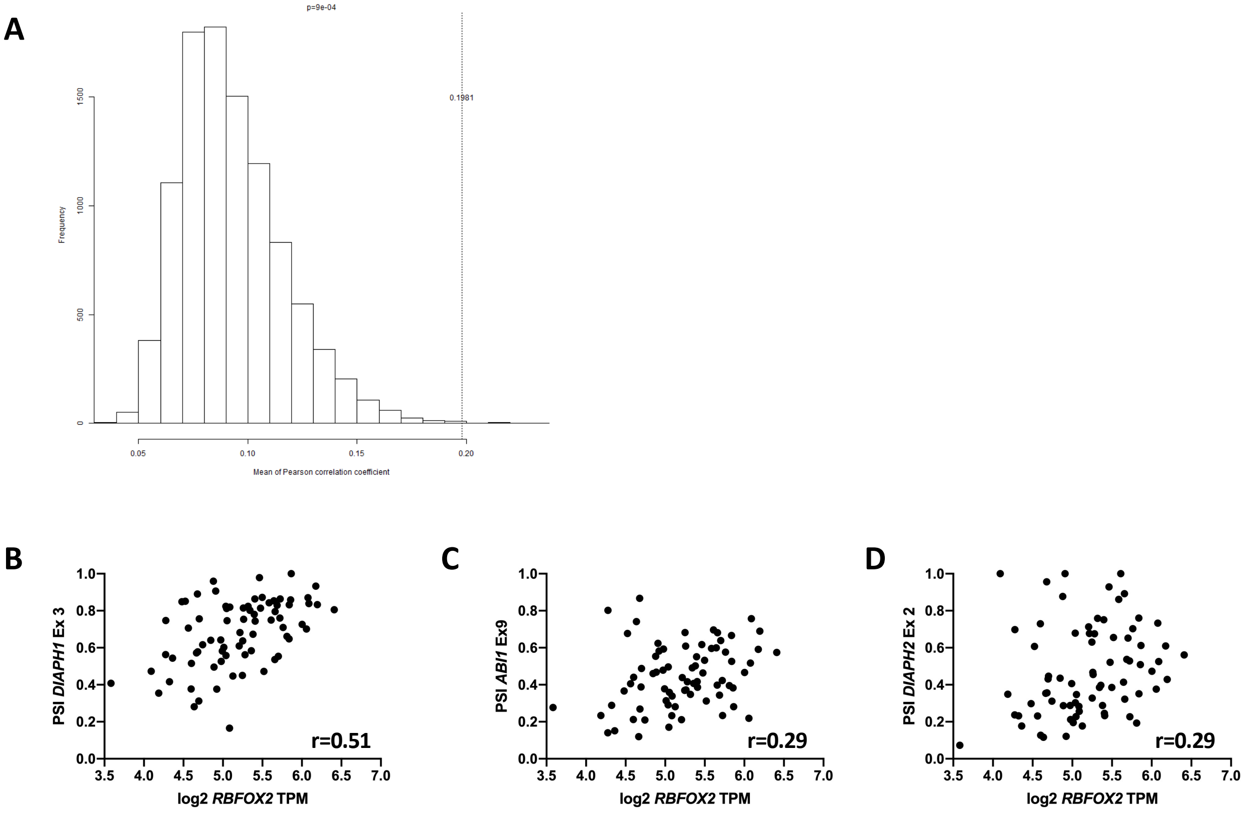

### Supplemental Figure 10

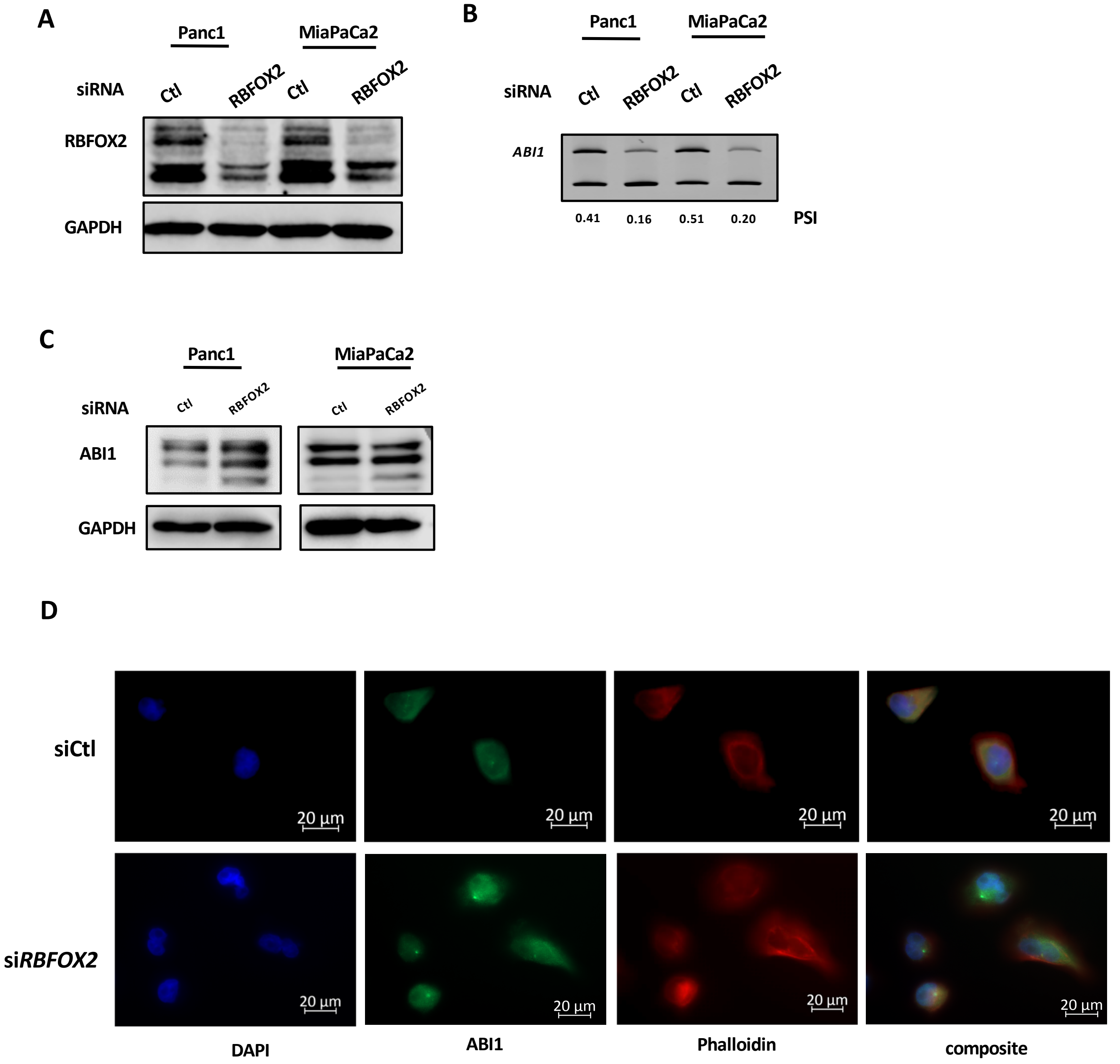
